# Transcriptomic response in symptomless roots of clubroot infected kohlrabi (*Brassica oleracea* var. *gongylodes*) mirrors resistant plants

**DOI:** 10.1101/391516

**Authors:** Stefan Ciaghi, Arne Schwelm, Sigrid Neuhauser

## Abstract

**Background:** Clubroot disease caused by *Plasmodiophora brassicae* (Phytomyxea, Rhizaria) is one of the economically most important diseases of *Brassica* crops. The formation of hypertrophied roots accompanied by altered metabolism and hormone homeostasis is typical for infected plants. Not all roots of infected plants show the same phenotypic changes. While some roots remain uninfected, others develop galls of diverse size. The aim of this study was to analyse and compare the intra-plant heterogeneity of *P. brassicae* root galls and symptomless roots of the same host plants (*Brassica oleracea* var. *gongylodes)* collected from a commercial field in Austria using transcriptome analyses.

**Results:** Transcriptomes were markedly different between symptomless roots and gall tissue. Symptomless roots showed transcriptomic traits previously described for resistant plants. Genes involved in host cell wall synthesis and reinforcement were up-regulated in symptomless roots indicating elevated tolerance against *P. brassicae*. By contrast, genes involved in cell wall degradation and modification processes like expansion were up-regulated in root galls. Hormone metabolism differed between symptomless roots and galls. Brassinosteroid-synthesis was down-regulated in root galls, whereas jasmonic acid synthesis was down-regulated in symptomless roots. Cytokinin metabolism and signalling were up-regulated in symptomless roots with the exception of one CKX6 homolog, which was strongly down-regulated. Salicylic acid (SA) mediated defence response was up-regulated in symptomless roots, compared with root gall tissue. This is probably caused by a secreted benzoic acid salicylic acid methyl transferase from the pathogen (PbBSMT), which was one of the highest expressed pathogen genes in gall tissue. The PbBSMT derived Methyl-SA potentially leads to increased pathogen tolerance in uninfected roots.

**Conclusions:** Infected and uninfected roots of clubroot infected plants showed transcriptomic differences similar to those previously described between clubroot resistant and susceptible hosts. The here described intra-plant heterogeneity suggests, that for a better understanding of clubroot disease targeted, spatial analyses of clubroot infected plants will be vital in understanding this economically important disease.

## Background

Clubroot disease is one of the most important diseases of *Brassica* crops worldwide accounting for approximately 10% loss in *Brassica* vegetable, fodder, and oilseed crops (Dixon, 2009). Clubroot is caused by *Plasmodiophora brassicae*, an obligate biotrophic protist, taxonomically belonging to Phytomyxea within the eukaryotic super-group Rhizaria (Bulman and Braselton, 2014; Neuhauser *et al.*, 2014). This soil borne pathogen has a complex life cycle: zoospores infect root hairs where primary plasmodia form. These plasmodia develop into secondary zoospores, which are released into the soil and re-infect the root cortex where secondary plasmodia develop (Kageyama and Asano, 2009). The secondary plasmodia mature into resting spores, which are released into the soil. In infected host tissue division and elongation of cells is triggered upon infection, which leads to hypertrophies of infected roots resulting in the typical root galls or clubroots (Fig. 1).

**Figure 1:**
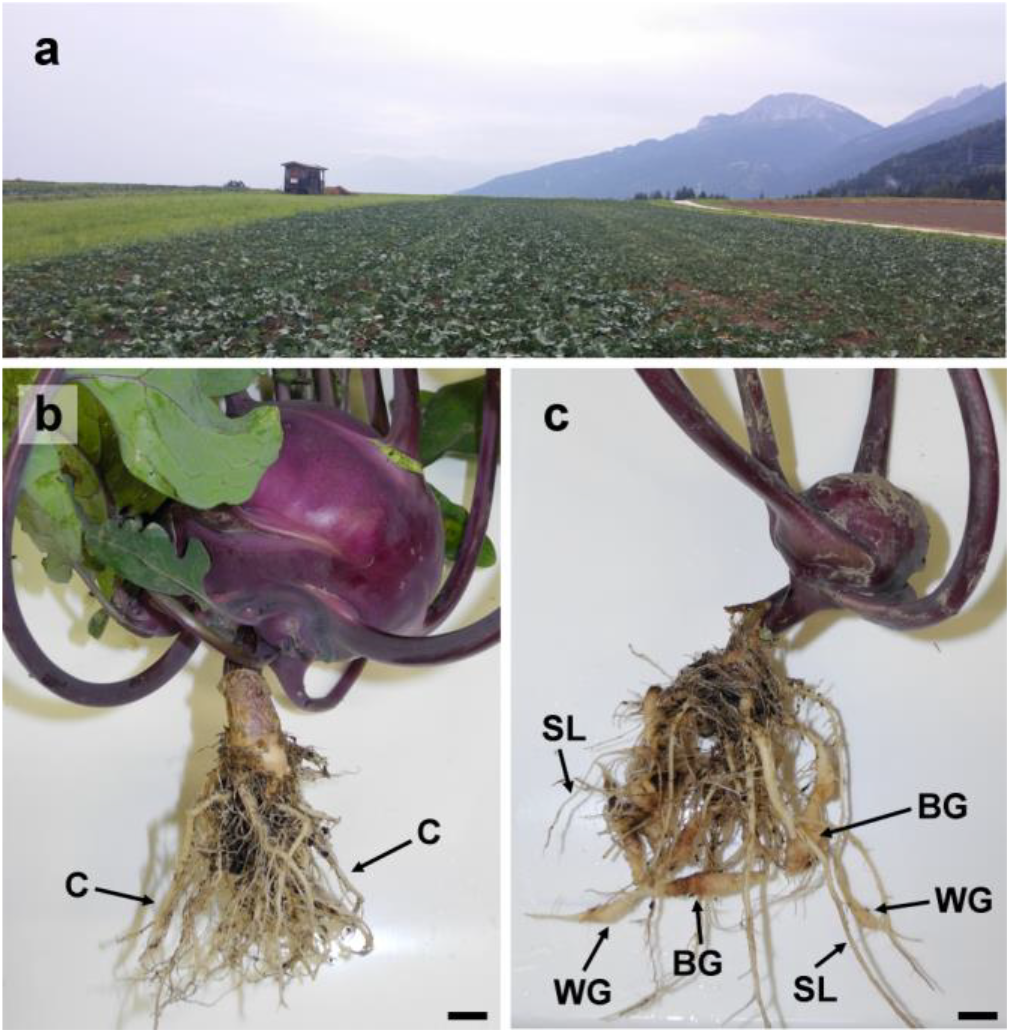
Sampling site and plant material. a: Field where plants were sampled (Ranggen, Tyrol; Austria). b: Normally developed kohlrabi plant. Roots were analysed as negative control (C). c: Clubroot infected kohlrabi plants with three different root phenotypes: symptomless roots (SL), white spindle galls (WG), and brownish spindle galls (BG). Scale bar: 1 cm.

*P. brassicae* can only be grown and studied in co-culture with its host. This has hampered both, targeted and large scale studies on the molecular basis of *P. brassicae* and the interactions with its host (Schwelm *et al.*, 2018). Because of the economic importance of clubroot disease, numerous studies analysed specific aspects of the biology, physiology and molecular biology of the plant-pathogen interaction to better understand and control the disease. The first of these experimental studies were based on the *Arabidopsis/Plasmodiophora* pathosystem (e.g. Devos *et al.*, 2006; Siemens *et al.*, 2006). During the last years an increasing number of *Brassica* (host) genomes became available (Wang *et al.*, 2011; Chalhoub *et al.*, 2014; Liu *et al.*, 2014; Cheng *et al.*, 2016), as did several *P. brassicae* genomes (Schwelm *et al.*, 2015; Bi *et al.*, 2016b; Rolfe *et al.*, 2016; Daval *et al.*, 2018) permitting new research approaches, including targeted transcriptome studies. There are analyses of (plant) transcriptomes of roots of clubroot infected plants compared with uninfected plants (Agarwal *et al.*, 2011; Bi *et al.*, 2016a; Malinowski *et al.*, 2016; Zhao *et al.*, 2017), of host varieties susceptible and tolerant to clubroot (Jubault *et al.*, 2008; Bi *et al.*, 2016a), or the host response to different *P. brassicae* isolates (Siemens *et al.*, 2006; Agarwal *et al.*, 2011; Lovelock *et al.*, 2013; Zhang *et al.*, 2016).

Plants infected with *P. brassicae* show marked physiological changes including cell wall biosynthesis, plant hormone metabolism and plant defence related processes. Expansin genes, involved in plant cell expansion and elongation (Cosgrove, 2005), were up-regulated in *P. brassicae* infected roots (Siemens *et al.*, 2006; Agarwal *et al.*, 2011; Irani *et al.*, 2018). In *P. brassicae* infected roots enzymatic activity of Xyloglucan endo Transglucosylase/Hydrolases (XTHs) increases (Devos *et al.*, 2005), while an early response to *P. brassicae* infection was up-regulation of the phenylpropanoid pathway that provides lignin precursors (Zhao *et al.*, 2017). With progression of clubroot development, the lignification of clubroot tissue was reduced (Deora *et al.*, 2013) and genes involved in lignification processes were down-regulated (Cao *et al.*, 2008). On the other hand cell wall thickening and lignification was suggested to limit the spread of the pathogen in tolerant *B. oleracea* (Donald *et al.*, 2008) and *B. rapa* (Takahashi *et al.*, 2001).

The development of clubroot symptoms is accompanied by changes of plant hormone homeostasis (Siemens *et al.*, 2006; Ludwig-Müller *et al.*, 2009; Malinowski *et al.*, 2016). During clubroot development, auxins mediate host cell divisions and elongation. Auxins increase over time during clubroot development and accumulate in *P. brassicae* infected tissues in a sink like manner (Ludwig-Müller *et al.*, 2009). In addition, genes belonging to the auxin conjugating GH3 protein family are regulated differentially during clubroot development (Jahn *et al.*, 2013) with one GH3 protein gene (PbGH3; CEP01995.1) identified in the *P. brassicae* genome (Schwelm *et al.*, 2015). Cytokinins (CKs) increase initially, but decrease again with the onset of gall formation (Malinowski *et al.*, 2016). At the same time *P. brassicae* plasmodia produce minute amounts of CKs (Müller and Hilgenberg, 1986). Therefore, CKs play a crucial role in disease development not only through their regulation of cell division but also through their interference in the sugar metabolism and invertase production, which might be crucial for the nutrition of *P. brassicae* (Siemens *et al.*, 2011; Walerowski *et al.*, 2018).

Stress- and defence related phytohormones like salicylic acid (SA), jasmonic acid (JA), brassinosteroids (BR), and ethylene (ET) and their regulatory pathways also change in response to pathogen infection (Kazan and Lyons, 2014). The accumulation of SA plays a key role in plant defence against biotrophic pathogens, often resulting in a localized hypersensitive response and induction of pathogenesis-related (PR) genes. Systemic acquired resistance (SAR) is a form of induced resistance that is activated by SA throughout a plant after being exposed to elicitors from microbes or chemical stimuli (Klessig *et al.*, 2018). High endogenous levels of SA and exogenous SA reduced the infection of the host by *P. brassicae* (Lovelock *et al.*, 2013; Lovelock *et al.*, 2016). In tolerant hosts SA related genes are induced upon infection (Siemens *et al.*, 2009; Agarwal *et al.*, 2011; Zhang *et al.*, 2016). The SAR-deficient *npr1-1* and SA-deficient isochorismate synthase 1 (ICS1) *sid2 Arabidopsis* mutants showed an increased susceptibility to *P. brassicae*, whereas the *bik-1* mutant, with elevated SA levels, was more resistant (Chen *et al.*, 2016b). Pathogenesis-related defence proteins were induced by SAR and expressed more highly in resistant than in susceptible *B. rapa* and *Arabidopsis* species (Jubault *et al.*, 2013; Bi *et al.*, 2016a; Jia *et al.*, 2017). *P. brassicae* might counteract the plant SA-mediated defence via a secreted methyltransferase (PbBSMT; AFK13134.1). This SABATH-like methyltransferase has been shown to convert SA to methyl-salicylate (MeSA) *in vitro* (Ludwig-Müller *et al.*, 2015). The proposed function *in planta* is the removal of SA in local infected tissue as MeSA is volatile. *Arabidopsis* mutants expressing the PbBMST gene showed a higher susceptibility towards *P. brassicae* (Bulman *et al.*, 2018).

The *P. brassicae* PbGH3 was also able to conjugate JA with amino acids *in vitro* (Schwelm *et al.*, 2015). In general, JA is associated with resistance against necrotrophic microbes (Pieterse *et al.*, 2012; Fu and Dong, 2013). In *A. thaliana* Col-0 several JA-responsive genes were induced in infected root tissues and JA accumulates in galls (Siemens *et al.*, 2006; Gravot *et al.*, 2012). *Jasmonate resistant 1* (*jar1*) mutants, impaired in JA-Ile accumulation, showed a higher susceptibility to *P. brassicae* (Agarwal *et al.*, 2011; Gravot *et al.*, 2012). Thus, JA responses contributed to a basal resistance against some strains of *P. brassicae* in *A. thaliana* Col-0 (Gravot *et al.*, 2012). But in partially resistant *Arabidopsis* Bur-0 only weak JA responses compared with the susceptible Col-0 were found (Lemarié *et al.*, 2015). Generally, clubroot susceptible hosts show a high level of JA response, whereas it is reduced in resistant hosts (Jubault *et al.*, 2008; Bi *et al.*, 2016a). Those differences might be due to occur if aliphatic or aromatic glucosinolate production is induced by JA in the particular host (Xu *et al.*, 2018).

The aim of this study was to generate the first data set of root tissue specific transcriptomic response of individual plants during clubroot development. Usually, clubroot infected plants do not develop symptoms uniformly on all root parts with some roots showing strong symptoms and others not showing symptoms at all (Fig. 1). We collected samples of kohlrabi (*Brassica oleracea* var. *gongylodes*) infected with *P. brassicae* (PbraAT) from a field in Austria. We compared morphologically different clubroots and symptomless roots of the same infected plants and a control plant. We analysed similarities and differences of their transcriptomic profile focussing on cell wall metabolism, hormone metabolism, and defence response.

## Methods

### Sampling

Kohlrabi “purple Vienna” plants with clubroots and without visible root infections were collected from a *P. brassicae* infested field in Ranggen (Tyrol, Austria, 47°15’27”N 11°12’37”E) in August 2016 with the consent of the farmer. No other permissions were necessary to collect these samples. Root samples were classified by visible properties into symptomless roots (SL), smaller white spindle galls with waxy appearance (WG) and larger brownish spindle galls (BG) (Fig. 1). Samples were taken in triplicates from three individual clubroot infected plants. Only one plant without apparent infection could be identified. All other plants collected from the vegetable plot without easily visible symptoms where infected when examined closer in the lab. Nevertheless we additionally took samples from the one uninfected control plant (C) (Fig. 1). Galls and roots were thoroughly washed with tap water before samples were taken (categories C, SL, WG, and BG) and transferred to RNA*later*^TM^ (Ambion, Austin, TX, USA) until RNA extraction.

### RNA extraction and sequencing

The outer layer of the root galls was trimmed-off and the trimmed galls and symptomless roots were snap-frozen in liquid nitrogen and transferred to 1.5 mL tubes containing RNase free zirconia beads (0.5 mm and 2 mm in diameter). Samples were homogenized using a FastPrep (MP Biomedicals, Santa Ana, CA, USA) for 40 s at 6 m s^−1^ followed by manual grinding with pestles after repeated snap-freezing. Total RNA was extracted using the Qiagen RNeasy Plant Mini Kit (Qiagen, Hilden, Germany) according to the manufacturer’s instructions, but with an additional 80% ethanol column wash prior elution. RNA quantity and quality were determined using Agilent Bioanalyzer 2100 (Agilent Technologies, Palo Alto, CA, USA). Additional RNA quality assessment, polyA selection (SENSE mRNA-Seq Library Prep Kit; Lexogen, Vienna, Austria), library construction (10 libraries; 1× C, 3× SL, 3× WG, and 3× BG) and sequencing was performed at the VBCF NGS Unit (Vienna, Austria). Sequencing was performed on the Illumina HiSeq 2500 platform (Illumina, San Diego, CA, USA) with a strand specific paired end library (2× 125 bp) using v4 chemistry.

### Bioinformatics

Raw reads were quality checked using FastQC (https://www.bioinformatics.babraham.ac.uk/projects/fastqc/). Illumina adapters were removed and good quality reads were kept using Trimmomatic v0.36 (sliding window 5 bp; average quality score > 20) (Bolger *et al.*, 2014). Only reads with a minimum length of 75 bp were processed further after a repeated FastQC check to confirm quality improvement. From the uninfected control library, 75% of the reads were randomly picked three times to generate pseudo-triplicates. Transcripts were *de novo* assembled using Trinity v2.2 (Grabherr *et al.*, 2011) with strand specific library type (RF) and jaccard clip options. Expression estimation was performed using Trinity embedded RSEM (Li and Dewey, 2011) keeping only transcripts with more than at least one fragment per kilobase per million (FPKM > 1) and an isoform percentage (IsoPct) > 1%.

The assembled transcripts were blasted using BlastN (Altschul *et al.*, 1990) against the coding sequences (CDS) of *B. oleracea* (Liu *et al.*, 2014; http://brassicadb.org/brad/datasets/pub/Genomes/Brassica_oleracea/V1.1/) and a custom database containing the CDS of the *P. brassicae* isolates e3 (Schwelm *et al.*, 2015) and PT3 (Rolfe *et al.*, 2016) to identify, if the transcript derived from the pathogen or host (E-value < 10^−5^). Transcripts with blast hits in both reference databases were analysed manually to identify their origin according to sequence identity and E-value. Transcripts with no hit in any reference were blasted (BlastP) against National Center for Biotechnology Information (NCBI) non redundant protein database and manually assigned to the corresponding species or discarded for further analysis. Transcripts with a best hit to a Brassicaceae reference sequence were assigned to the host transcriptome. Transcripts with hits to *P. brassicae* sequences were assigned to the pathogen transcriptome. Open reading frames (ORFs) were predicted using TransDecoder v3.0.1 (https://github.com/TransDecoder/). Only the longest ORF per transcript was used for further analysis. Translated peptide sequences were annotated using the KEGG (Kyoto Encyclopedia of Genes and Genomes) Automatic Annotation Server (KAAS) (Moriya *et al.*, 2007) and eggNOG-mapper v0.99.3 (Huerta-Cepas *et al.*, 2016). Kohlrabi genes were additionally annotated using Mercator (Lohse *et al.*, 2014) with default settings. Mercator categories were used for MapMan v3.6.0RC1 (Usadel *et al.*, 2009) to bin predicted genes into groups. Putative secreted proteins of *P. brassicae* were predicted with Phobius v1.01 (Käll *et al.*, 2004) and SignalP v4.1 (Nielsen, 2017) in combination with TMHMM v2.0 (Krogh *et al.*, 2001). Carbohydrate active enzymes were predicted using dbCAN (Yin *et al.*, 2012).

Log_2_-fold changes of differentially expressed genes (DEGs) were calculated using edgeR with default settings (Robinson *et al.*, 2010). All DEGs with false discovery rate (FDR) < 0.05 were treated as DEGs. Heatmaps for selected Mercator/MapMan categories were created using R v3.3.2 (R Core Team, 2016) with the package ‘pheatmap’ v1.0.8 (https://cran.r-project.org/web/packages/pheatmap/index.html) applying UPGMA clustering. Labelling of the predicted *B. oleracea* genes was done according to their homologous *A. thaliana* genes from TAIR (The Arabidopsis Information Resource) and adapted if necessary. Abundance of DEGs was visualized using the R package ‘ggplot2’ v2.2.1 (Wickham, 2009).

## Results

### Transcriptome analyses

In total 162 million good quality reads with an average length of 125 bp were obtained from all libraries (Additional file 2: Table S1). A total of 10940 genes including isoforms were predicted for *P. brassicae* and 42712 for *B. oleracea*. About 50% of the *P. brassicae* and 85% of the kohlrabi transcripts could be functionally annotated using eggNOG-mapper. Only 0.0005% of the reads of the control plant (C) and the symptomless root samples (SL) matched *P. brassicae* transcripts, which indicates that these roots were not infected by *P. brassicae*. In the white spindle galls (WG) and brownish spindle galls (BG) libraries 23% and 33% of the reads matched *P. brassicae* (Additional file 1: Figure S1).

The transcriptomes of infected plants (SL, WG, BG) contained a total of 5204 differentially expressed genes (DEGs). Compared with SL, in WG 1619 DEGs were up-regulated and 2280 were down-regulated (Fig. 2, Additional file 2: Table S2), while in BG 942 DEGs were up- and 2571 were down-regulated. 790 plant DEGs were assigned to the COG (Cluster of Orthologous Groups) category “Information and Storage Processing”, 1401 to “Metabolism”, 1245 to “Cellular Process and Signalling” and 1768 to “Poorly Characterized” by eggNOG-mapper (Fig. 2, Additional file 2: Table S3). Only 19 plant genes were differentially expressed between BG and WG (Additional file 2: Tables S4). Between the control plant and the three root tissue types of the infected plants (C vs SL/WG/BG) 19230 DEGs were found (Additional file 2: Tables S5, S6).

**Figure 2:**
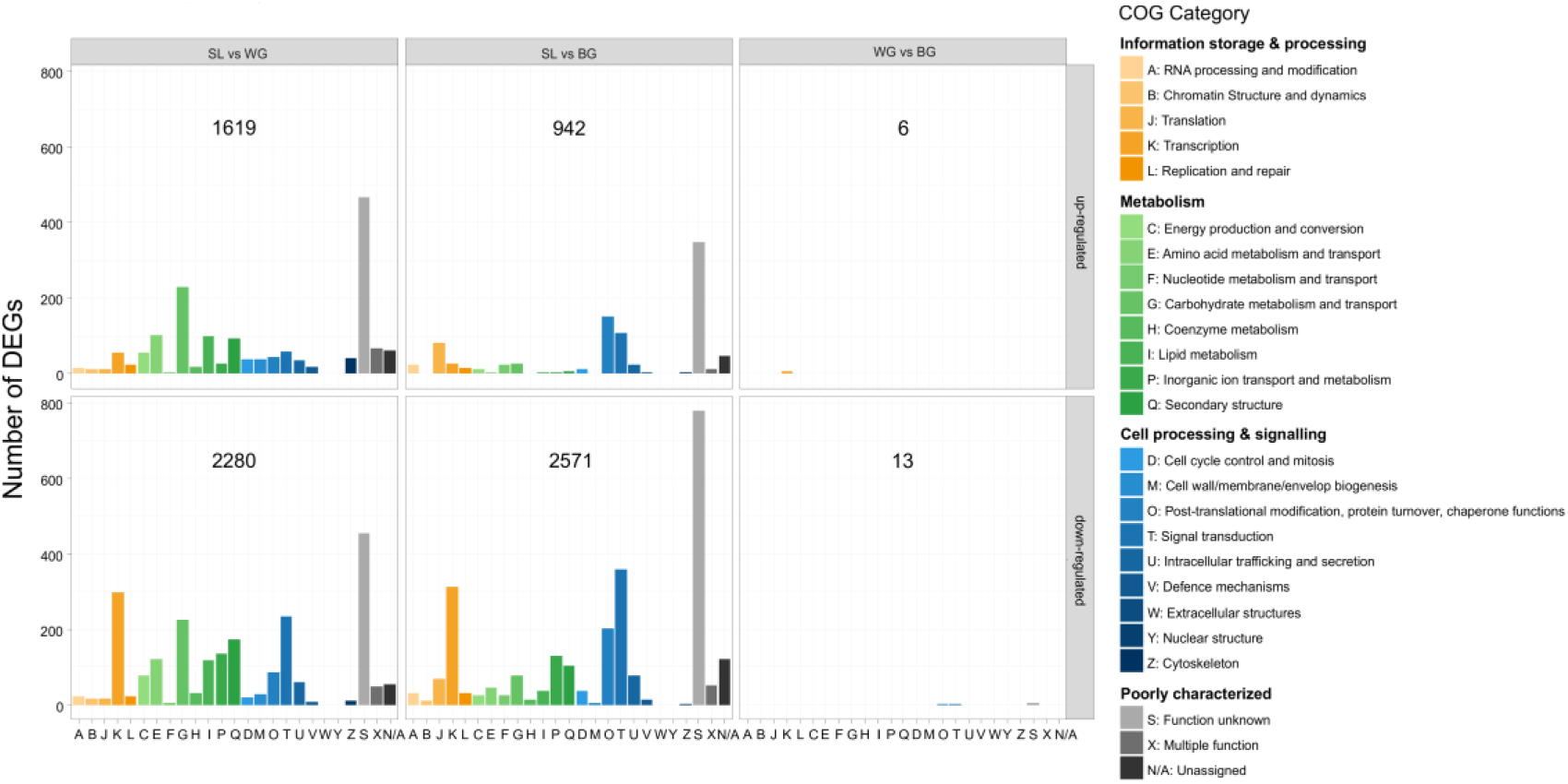
Numbers of differentially expressed genes (DEGs) in clubroot infected kohlrabi roots per COG category. DEGs were split into up- and down-regulated genes. Total number of DEGs in each panel is given.

### Plant cell wall metabolism

In *B. oleracea* 161 of the 5204 DEGs within infected plants (SL vs WG vs BG) were involved in cell wall synthesis, modification, degradation, or phenylpropanoid metabolism. Cellulose, hemicellulose, pectin, and lignin synthesis genes involved in cell wall reinforcement were up-regulated in SL compared with gall tissues (Fig. 3), whereas genes involved in cell wall modification and degradation were down-regulated (Fig. 3). The changes in expression of cell wall genes were more prominent between SL and BG than between SL and WG. In SL a UDP-D-glucuronate 4-epimerase 6 (GAE6) homolog responsible for the synthesis of UDP-D-glucuronic acid, the main building block for pectins (Harholt *et al.*, 2010) was up-regulated compared with gall tissue. Predicted expansin (EXP) and expansin-like (EXL) genes were mainly down-regulated in SL compared with WG and BG (Fig. 3). Genes coding for XTHs were among the strongest down-regulated DEGs in SL, with XTH24 being the strongest down-regulated transcript of all DEGs. The phenylpropanoid pathway was up-regulated in SL compared with WG and BG (Fig. 3). This includes the phenylalanine ammonia-lyase 1 (PAL1) homolog, a key enzyme in lignin biosynthesis. Lignin is also part of the xylem and xylogenesis genes which were up-regulated in SL compared with BG. Flavonoid metabolism was also induced in SL (Additional file 1: Figure S2). Compared with an uninfected control plant cell wall synthesis genes were up-regulated in SL (Additional file 1: Figure S3).

**Figure 3:**
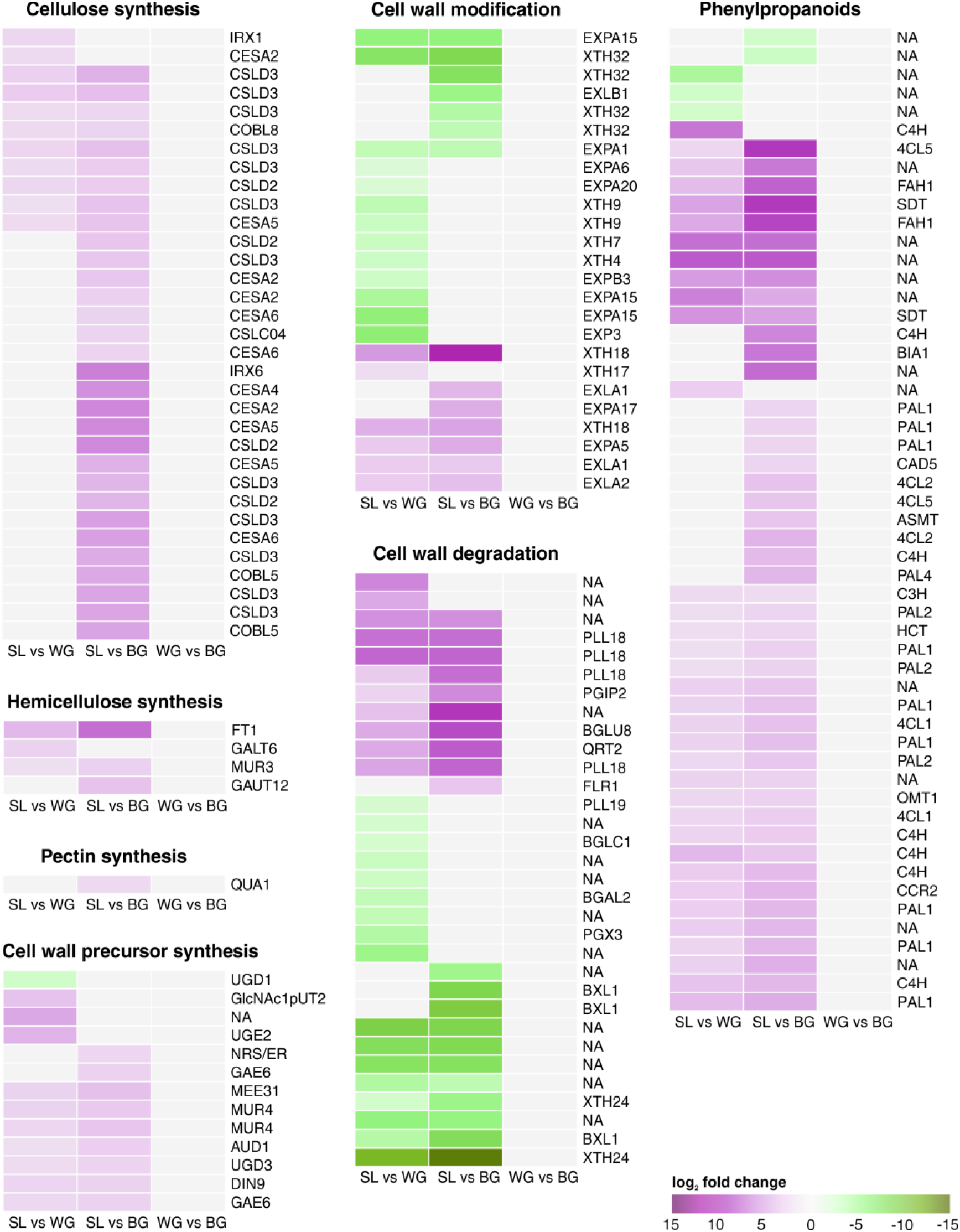
Kohlrabi cell wall metabolism. Clustered heatmaps of log_2_ fold change values of DEGs. Genes involved in anabolic processes of cell wall components were generally up-regulated in SL compared with WG and BG, whereas catabolic and modifying genes were mainly down-regulated. No DEGs were present comparing WG with BG. Up-regulated genes are shaded in purple and down-regulated genes in green. *Arabidopsis* homologs are given. Genes are annotated and categorized according to MapMan/Mercator. NA: not assigned by MapMan. Supplementary information for the genes including KEGG, eggNog, and TAIR annotation are given in Additional file 3.

### Plant hormones

The CK and auxin metabolism was altered in WG and BG compared with SL (Fig. 4). So was a homolog of CKX6 (cytokinin oxidase/dehydrogenase 6) down-regulated in SL compared with WG. Homologs of CKX5, CK receptors, and a CK-regulated UDP-glucosyltransferase were up-regulated in SL. The CK synthesis genes CYP735A2 (cytochrome P450) and LOG1 (lonely guy 1) were up-regulated in SL compared with root galls (Fig. 4). Down-stream in CK-signalling we found DEGs within ARR (Arabidopsis response regulator; CK-signalling target) genes (Fig. 4). Type-B ARRs were differentially expressed in WG and BG whereas differentially expressed Type-A ARRs were exclusively found in BG. A putative AHK4 (Arabidopsis histidine kinase 4; CK-receptor) gene was up-regulated in SL compared with BG where it was not detected although expression values in SL were low. No DEGs of AHPs (Arabidopsis histidine phosphotransfer protein) were found in SL compared with root galls. Most auxin related DEGs, auxin response factors (ARFs and IAAs) and IAA amino acid conjugate synthetases (GH3) were up-regulated in SL compared with gall tissue (Additional file 1: Figure S4). However, an IAA7 (repressor of auxin inducible genes) and a homolog of GH3.2 were down-regulated. Expression of PIN-FORMED 1 (PIN1) genes was reduced in SL (Fig. 4), whereas SAUR (small auxin up-regulated RNA) and AIR12 (auxin-induced in root cultures protein 12-like) genes, were found in up- and down-regulated DEGs (Fig. 4). Myrosinases and nitrilases were up-regulated in SL compared with galls (Additional file 1: Figure S5). Compared with the unifected control both CK and auxin metabolism were up-regulated in SL (Additional file 1: Figure S3). In BG two transcripts related to auxin synthesis and regulation were down-regulated compared with WG (Additional file 2: Table S4).

**Figure 4:**
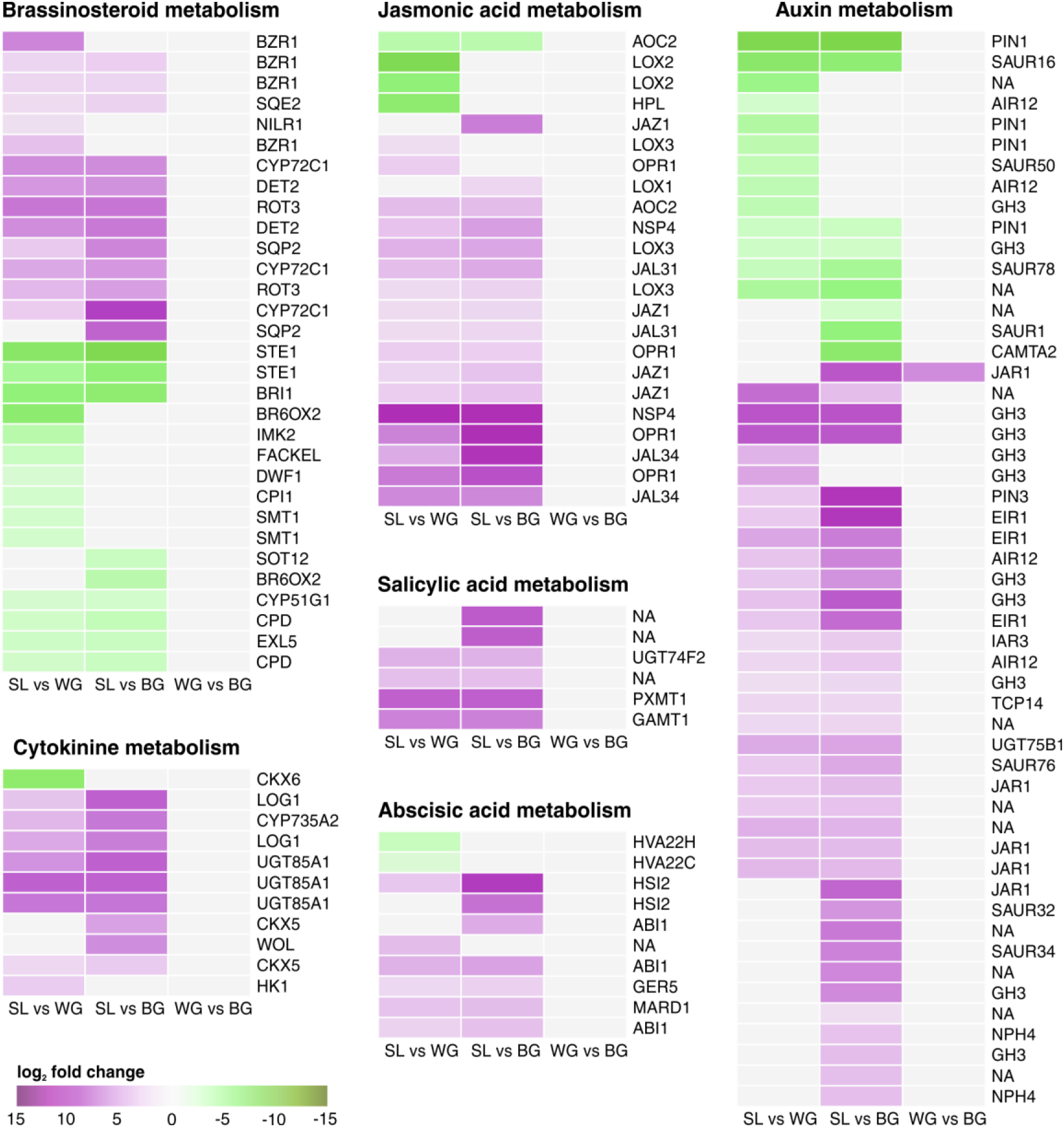
Kohlrabi phytohormone metabolism. Heatmaps of log_2_ fold change values of DEGs. Genes involved in cytokinin, jasmonic acid, salicylic acid, and abscisic acid metabolism were up-regulated in SL. Genes coding for brassinosteroids clustered into two groups: later stages in BR biosynthesis (down-regulated) and early sterol biosynthesis (up-regulated). Genes involved in auxin metabolism were found within the up- and down-regulated DEGs in SL compared with root galls. One DEGs (JAR1) was found between WG and BG. Up-regulated genes are shaded in purple and down-regulated genes in green. *Arabidopsis* homologs are given. Genes are annotated and categorized according to MapMan/Mercator, except for cytokinin metabolism for which genes were categorized as described by (Malinowski *et al.*, 2016). NA: not assigned by MapMan. Supplementary information for the genes including KEGG, eggNog, and TAIR annotation are given in Additional file 4.

Early sterol biosynthesis genes, such as the steroid reductase DET2, were up-regulated in SL compared with galls (Fig. 4). However, BR biosynthesis genes were generally down-regulated in SL (Fig. 4), including key genes like DWF1 (dwarf 1) or BRI1 (BR receptor brassinosteroid insensitive 1).

Abscisic acid (ABA) signal transduction related genes like ABI1 (ABA insensitive 1) and HSI2 (high-level expression of sugar-inducible gene 2) were up-regulated in SL compared with WG and BG (Fig. 4) whereas ABA related transcription factors WRKY18 and HVA22A/C homologs were down-regulated in SL (Fig. 4, Additional file 1: Figure S6).

### Plant defence

Generally, genes for disease resistance proteins were up-regulated in SL compared with WG and BG (Fig. 5, Additional file 1: Figure S7). From pathogen recognition genes via signalling proteins and transcription factors to pathogenesis related (PR) proteins, the whole signal cascade of pathogen defence was affected. Of the predicted defence related DEGs within infected plants (SL vs WG vs BG) 60 were assigned as TIR-NBS-LRR (Toll/interleukin-1 receptor nucleotide-binding site leucine-rich repeat) class proteins. Within R-genes only PP2-A (phloem protein 2-A) genes were up-regulated in SL compared with root galls (Fig 5). Expressed genes for disease resistance protein RSP4 were not altered between the tissue types (Additional file 5). Signalling genes encoding for MLO (mildew resistance locus o) and MKS1 (MAP kinase substrate 1) were up-regulated in SL (Fig. 5), while no differences in the expression of CHORD or SRFR1 (suppressor of RSP4-RDL1) genes was observed (Additional file 5). Two transcription factors involved in the biotic stress signalling cascade (MapMan bin 20.1.5) were up-regulated in SL compared with the root galls (Fig 5). MAP3K (mitogen activated kinase kinase kinase) and RSH (RELA/SPOT homolog) genes were not differentially expressed within infected plants (Additional file 5).

**Figure 5:**
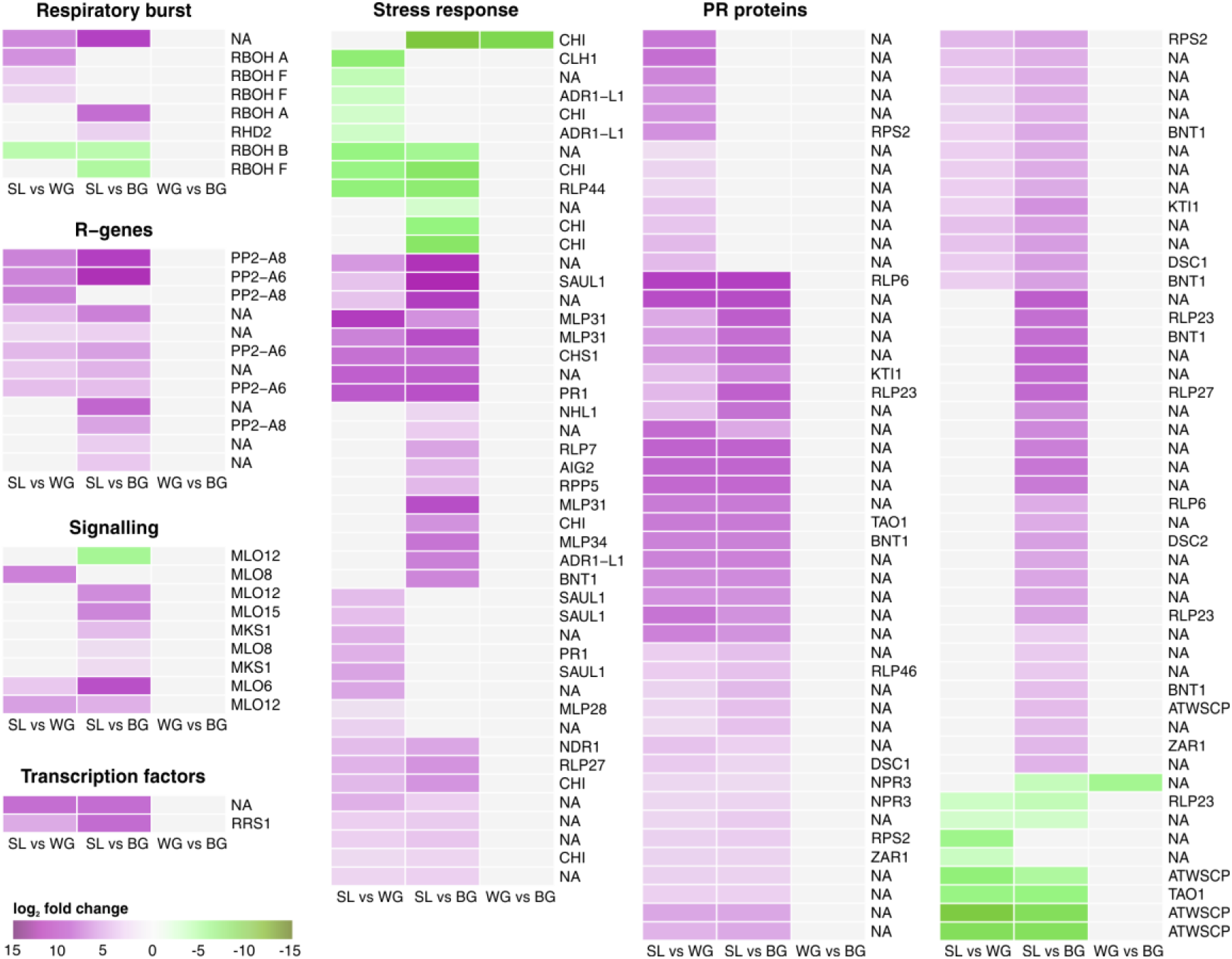
Biotic stress response of kohlrabi. Heatmaps of log_2_ fold change values of DEGs. Almost all DEGs were up-regulated in SL compared with root galls. Two down-regulated DEGs, a chitinase (CHI) and a gene of unknown function, were found between WG and BG. Up-regulated genes are shaded in purple and down-regulated genes in green. *Arabidopsis* homologs are given. Genes are annotated and categorized according to MapMan/Mercator. NA: not assigned by MapMman. Supplementary information for the genes including KEGG, eggNog, and TAIR annotation are given in Additional file 5.

The BIK1 (botrytis-induced kinase 1), which interacts with BRI1 and BAK1 (BRI1-associated receptor kinase) to induce defence responses was up-regulated in SL compared with galls (Additional file 1: Figure S8) and up-regulated in SL compared with C (Additional file 1: Figure S3).

In the SL samples JA-related genes such as LOX2 (lipoxygenase 2), AOC (allene oxide cyclase), and HPL (hyperoxide lyase) were down-regulated, while other LOX genes and the JA amido synthetase genes JAR1 were up-regulated in SL compared with galls (Fig. 4). One down-regulated isoform of JAR1 was found between BG and WG. We found no glucosinolate biosynthesis genes in the DEGs in our study.

The ICS1 gene and the ICS1 activating transcription factor WRKY28 (van Verk *et al.*, 2011) were up-regulated in SL compared with the control plant (data not shown). Genes for SA modification, like the SA methylating SABATH methyl transferase genes (PXMT1, GAMT1) and a SA-glucosidase (UGT74F2) were up-regulated in SL compared with galls (Fig. 4). The SA induced PR1 gene was induced in SL (Fig. 5, Additional file 1: Figure S7). The PR-gene expression regulator NPR1 (non-expressor of PR1) was not differentially expressed in our samples. Of genes that regulate PR1 expression via NPR1, WRKY70 was down-regulated in SL whereas NPR3 and TGA3 were up-regulated (Fig. 5, Additional file 1: Figure S7). Compared with the uninfected plant, the TGA3 gene expressions appeared to be induced in SL. Genes for the TAO1 disease resistant protein, which induces PR1 expression were up-regulated in SL, as well as the NDR1 (non-race specific disease resistance 1) gene, required for the establishment of hypersensitive response and SAR. Genes for the disease resistant protein RPS2, activated by NDR1, were also up-regulated in SL (Fig. 5, Additional file 1: Figure S7). Additionally, other defence related genes coding for protease inhibitor genes, R-genes or some chitinases were down-regulated in SL (Fig. 5). Comparing biotic stress response genes of SL with the control plant (C), we observed an up-regulation of 164 of the total 190 DEGs in SL (Additional file 1: Figure S3).

### *P. brassicae* gene expression

The *P. brassicae* genes with the highest FPKM values belonged to growth and cellular process related COG categories such as translation, transcription, and signal transduction, but also in energy conversion and carbohydrate and lipid metabolism (Additional file 1: Figure S9). The *P. brassicae* PbBSMT gene was amongst the highest expressed pathogen genes (Additional file 2: Tables S7, S8). Other highly expressed genes were HSPs (heat shock proteins), a glutathione-S-transferase, an ankyrin repeat domain-containing protein, ribosomal genes, and genes of unknown function (Additional file 2: Tables S7, S8). The PbGH3 gene was not expressed in our samples. The *P. brassicae* protease gene PRO1, proposed to be involved in spore germination (Feng *et al.*, 2010), was expressed in both WG and BG.

Between WG and BG samples only five *P. brassicae* DEGs were identified, coding for a HSP, a chromosomal maintenance protein, a DNA-directed RNA-polymerase, a retrotransposon and a Scl Tal1 interrupting locus protein (Additional file 2: Table S9). Cumulating all FPKM values revealed that most sequenced reads from *P. brassicae* RNA extracted from root galls (WG and BG) mapped to the COG categories “Post-translational modification, protein turnover, chaperon functions” and “Translation” (Additional file 1: Figure S10). Very few *P. brassicae* reads were found in the data obtained from the control plant and SL (Additional file 1: Figure S11), those were most likely from attached spores or contamination via soil particles.

## Discussion

### Symptomless roots of clubroot infected plants show transcriptomic traits of clubroot resistant/tolerant plants

We found that symptomless roots and clubroots originating from the same plant showed differences in their transcriptomic profile similar to previously described differences of roots between resistant and susceptible plants (Chen *et al.*, 2016a; Jia *et al.*, 2017). Gene expression patterns of symptomless roots were similar to the patterns described for resistant hosts, while in clubroot tissue patterns were similar to those observed in susceptible plants.

Reinforcement of cell walls has previously been reported to hamper the development of *P. brassicae* in resistant *B. oleracea* (Donald *et al.*, 2008) and *B. rapa* callus cultures (Takahashi *et al.*, 2001). Lignin biosynthesis genes were up-regulated in SL tissue compared with root gall tissue (Fig. 3). Induced lignification processes were observed in shoots of infected plants (Irani *et al.*, 2018) and between resistant and susceptible *B. oleracea* cultivars (Zhang *et al.*, 2016). PAL1, a key enzyme in lignin, SA (discussed below) and flavonoid biosynthesis (Chu *et al.*, 2014; Song *et al.*, 2016; Lahlali *et al.*, 2017) was up-regulated in symptomless roots compared with clubroot tissue (Fig. 3). Increased lignin biosynthesis and up-regulation of PAL1 has been described for a clubroot resistant oilseed *B. rapa* line carrying the resistance gene Rcr1 (Lahlali *et al.*, 2017), while callus cultures overexpressing PAL1 were resistant to infection by *P. brassicae* (Takahashi *et al.*, 2001). Root reinforcement, therefore, seems to be a part of the tolerance mechanisms of plants against *P. brassicae*, which has only a limited arsenal of plant cell wall degrading enzymes in its genome (Schwelm *et al.*, 2015; Rolfe *et al.*, 2016) and infects its hosts via mechanical force with a specialised extrusosome called “Stachel and Rohr” (Kageyama and Asano, 2009). Once *P. brassicae* successfully infected its host, movement and spread within root tissues has been discussed to happen via the plasmodesmata (Donald *et al.*, 2008; Riascos *et al.*, 2011; Badstoeber *et al.*, 2018). Increased stability of the cell wall, as indicated by gene expression patterns in symptomless roots (Fig. 3) or described for resistant plants (Takahashi *et al.*, 2001; Donald *et al.*, 2008), requires higher mechanical force for successful (primary) infection of yet uninfected roots or movement in between host cells. Therefore, it is likely that cell wall reinforcement of the host cells is a considerable obstacle for *P. brassicae*. The result of increased lignification in SL compared with galls could also be the result of repressed xylem development in *P. brassicae* infected hosts (Malinowski *et al.*, 2012).

In symptomless roots of infected plants, defence related pathways regulated by hormones show patterns usually linked to induced plant defence (Fig. 4). Clubroot tissue on the other hand showed suppression of SA-defence related processes. Genes of SA biosynthesis were up-regulated in SL, but were down-regulated in galls. Salicylic acid can be synthesised via isochorismate (ICS pathway) or from phenylalanine (via PAL pathway) (Pieterse *et al.*, 2012; Lovelock *et al.*, 2016). The up-regulation of the PAL1 gene in symptomless roots could therefore also be linked to the PAL-dependent synthesis of SA, but the majority of SA in clubroot is produced via the ICS pathway (Lovelock *et al.*, 2016). Based on the higher expression of ICS1 and WRKY28 in SL compared with the control, the synthesis of SA was likely induced in SL. In clubroot resistant hosts SA-defence is usually induced (Jubault *et al.*, 2013; Bi *et al.*, 2016a; Lovelock *et al.*, 2016), whereas the SA deficient *sid2* (ICS1) mutant of *Arabidopsis* was more susceptible to *P. brassicae* infection (Chen *et al.*, 2016b).

High SA levels in plant tissues helped to reduce new *P. brassicae* infections (Lovelock *et al.*, 2013), but SA alone is not sufficient to induce resistance against *P. brassicae* (Lovelock *et al.*, 2016). Because SA levels increase in clubroot tissue *P. brassicae* is thought to secret a SABATH-type methyltransferase (PbBSMT; Ludwig-Müller *et al.*, 2015; Bulman *et al.*, 2018), which was one of the highest expressed genes of *P. brassicae* in this study (Additional file 2: Tables S7, S8). PbBSMT has been shown to methylate salicylic acid, contributing to a local reduction of SA in the galls (Ludwig-Müller *et al.*, 2015). MeSA is the major transport form of SA in the plant and has a key role in inducing SAR (Park *et al.*, 2007; Vlot *et al.*, 2009). So based on our data we hypothesise that *P brassicae* reduces SA concentrations in the galls via PbBMST mediated methylation (Ludwig-Müller *et al.*, 2015). The produced MeSA could trigger SA-related defences in distant plant parts, which become resilient towards new pathogen infection.

Processes downstream from SA are mediated via NPR1 (not differentially expressed in our dataset). NPR1 induces PR-gene expression when bound to TGA3 (Saleh *et al.*, 2015). In its interaction with WRKY70, NPR1 serves as a negative regulator of the SA biosynthesis gene ICS1 (Wang *et al.*, 2006). Thus the observed up-regulation of TGA3 in SL (Additional file 1: Figure S7) would lead to an induction of expression of PR-genes in SL. The reduced expression of WRKY70 in SL compared with the galls would lead to a higher SA production in the uninfected root tissue. Additionally, we found an up-regulation of NPR3 genes, coding for repressors of SA-defence genes (Ding *et al.*, 2018), in SL compared with the galls (Fig. 5). When bound to SA, NPR3 would lose its function of repressing SA-defence genes (Ding *et al.*, 2018). The higher expression of NPR3 in the SL might be necessary to compensate for negative effects of the SA synthesis in SL roots.

Jasmonic acid contributed to a basal resistance against some strains of *P. brassicae. A. thaliana* Col-0 and *A. thaliana* mutants impaired in JA-Ile accumulation showed a higher susceptibility to *P. brassicae* (Agarwal *et al.*, 2011; Gravot *et al.*, 2012). In *A. thaliana* Col-0 several JA-responsive genes were induced in infected root tissues and JA accumulates in galls (Siemens *et al.*, 2006; Gravot *et al.*, 2012). But in partially resistant *Arabidopsis* Bur-0 only weak JA responses compared with the susceptible Col-0 were found (Lemarié *et al.*, 2015). Those differences might be due to if aliphatic or aromatic glucosinolate production is induced by JA in the particular host (Xu *et al.*, 2018). Generally, clubroot susceptible hosts show a high level of JA response, whereas it is reduced in resistant hosts (Jubault *et al.*, 2008; Bi *et al.*, 2016a). In our samples, JAZ genes were up-regulated in SL compared with the gall tissue (Fig. 4). JAZ proteins act as JA co-receptors and transcriptional repressors in JA signalling (Kazan and Manners, 2012) reducing the JA synthesis in SL. This was previously observed in *B. oleracea* plants, where JAZ expression was up-regulated in resistant plants and JA synthesis was highly induced in susceptible plants (Zhang *et al.*, 2016). Among JA-metabolism genes HPL, and LOX2 expression were up-regulated in the galls. LOX2 is essential for the formation of oxylipin volatiles (Mochizuki *et al.*, 2016). The HPL protein competes with substrates essential for JA-synthesis, producing volatile and non-volatile oxylipins (Wasternack and Hause, 2013). The higher expression of HPL and LOX2 in the galls might lead to the production of volatile aldehydes rather than a JA accumulation in the galls.

In SL BR-synthesis and BR-signalling genes were down-regulated (Fig. 4). BRs are necessary for the development of clubroot tissue (Schuller *et al.*, 2014), hence a reduction in BRs impairs *P. brassicae* growth and development in SL. The receptor-like cytoplasmic kinase BIK1, a negative regulator of BR-synthesis (Lin *et al.*, 2013), was induced in SL (Additional file 1: Figure S8). *Arabidopsis bik1* knockout mutants have an increased tolerance to *P. brassicae* lacking the typical pathogen phenotype (Bi *et al.*, 2016a; Chen *et al.*, 2016c). For BIK1 the expression of the SL roots differed to resistant plants, as *Arabidopsis bik1* knockout mutants have an increased tolerance to *P. brassicae* (Bi *et al.*, 2016a; Chen *et al.*, 2016c). In *Arabidopsis bik1* clubroot resistance was likely not due to the regulatory function of BIK1 for BR, but to increased PR1 expression in this mutant (Chen *et al.*, 2016c).

### Transcriptomes of symptomless roots and clubroots of the same plant are markedly different

Clubroots and symptomless roots of the same *P. brassicae* infected plants showed markedly different gene expression patterns. The morphological changes of the clubroots go in hand with expression of genes that reduce cell wall stability and growth related processes. Genes for cell wall-loosening processes such as expansins and XTHs (Downes *et al.*, 2001; Sun *et al.*, 2005), were up-regulated in gall tissue (Fig. 3). Up-regulation of expansins was reported from *Arabidopsis* clubroot tissue (Siemens *et al.*, 2006; Irani *et al.*, 2018) while XTH activity was reported in *B. rapa* clubroots (Devos *et al.*, 2005). Suppression of xyloglucan, xylan, hemicellulose, pectin and lignin synthesis in gall tissues implies additional reduction of the cell wall stability and rigidity in gall tissues similar to previous findings (Schuller *et al.*, 2014; Zhang *et al.*, 2016). GAE6 expression was reduced in the galls supporting clubroot development: *Arabidopsis gae1*, *gae6*, and *gae1/gae6* mutants contained lower levels of pectin in their leaf cell walls making them more susceptible to *Pseudomonas syringae* and *Botrytis cinerea* (Bethke *et al.*, 2016). A similar mechanism might benefit *P. brassicae*. Induction of the lignin pathway was an early response (48h) in *Arabidopsis* (Zhao *et al.*, 2017). In the brownish kohlrabi galls investigated in our study, lignification genes were down-regulated suggesting *P. brassicae* suppresses host lignification in tissues where it is already established. In *B. napus* reduced lignification was also implied by the down-regulation of CCoAOMT (caffeoyl-CoA O-methyltransferase) (Cao *et al.*, 2008). Here, although not statistically significant, CCoAOMT was also lower expressed in root galls compared with SL, implying a decreased lignin biosynthesis in clubroots.

The marked hypertrophies of the plant roots go hand in hand with changes in the homeostasis of the growth hormones CK and auxins which appear to be host and time dependent (Ludwig-Müller *et al.*, 2009; Jia *et al.*, 2017). With the exception of CKX6, cytokinin related genes were up-regulated in SL (Fig. 4). Fine tuning of the hormone balance appears to be essential in clubroot disease development (Ludwig-Müller *et al.*, 2009). Elevated CK levels are important for the onset of disease development via increasing cell divisions. However, at the onset of gall formation, CK metabolism genes including CK synthesizing and degrading enzymes are repressed (Siemens *et al.*, 2006; Malinowski *et al.*, 2016). The more active CK metabolism in the SL might interfere with clubroot development as CKX overexpressing *Arabidopsis* mutant showed reduced gall formation (Siemens *et al.*, 2006). In our kohlrabi samples, CKX6 was strongly down-regulated in SL compared with WG but not compared with BG (Fig. 4). The high expression of CKX6 in WG indicates the presence of plasmodia, as the gene was found to be strongly up-regulated only in cells containing *P. brassicae* plasmodia (Schuller *et al.*, 2014). Although AHK4 was described to be highly induced in infected *A. thalinana* plants 10 dpi (Siemens *et al.*, 2006), in a different study expression of AHK and AHP genes did not differ between infected and uninfected *Arabidopsis* at later time points (Malinowski *et al.*, 2016). Similar to these findings, in our much longer infected kohlrabi plants AHK and AHP genes were not differential expressed between SL and root galls. Expression levels were also very low indicating no major role of those genes in the analysed roots. The ARR5 gene was down-regulated in infected roots and hypocotyl tissues 16 and 26 dpi in *A. thaliana* (Malinowski *et al.*, 2016) but in a different study ARR5 was up-regulated 10 dpi in infected roots (Siemens *et al.*, 2006). For *Brassica* hosts, the expression of ARR5 in clubroot infected roots has not been described. In our samples ARR5 levels increase in BG, but no difference is seen between SL and WG. The ARR5 gene does not appearto have a prominent role at the stage of the here investigated infections. Also. evidence that *P. brassicae* interferes with the CK balance via the PbCKX of *P. brassicae* in gall tissues has not been seen in this study as the PbCKX gene was not expressed.

Myrosinases and nitrilases that can synthesize auxins from secondary metabolites or aromatic amino acids were up-regulated in SL compared with galls (Additional file 1: Figures S6), but were previously reported to be induced in *Arabidopsis* galls (Grsic-Rausch *et al.*, 2000; Siemens *et al.*, 2006). The auxin-induced GH3 gene family, which conjugates IAA to several amino acids, is involved in various responses of plants to abiotic and biotic stresses. The GH3.2 gene was shown to be specifically expressed in clubroots of *Arabidopsis* (Jahn *et al.*, 2013), and was also up-regulated in the kohlrabi galls. Expression of the *P. brassicae* PbGH3 gene was not detected in this study and appears not to play a role in the auxin (or JA) homeostasis at the stage of our gall samples.

Besides the described defence responses in SL that are similar to defence responses of resistance plants, we observed up-regulation of other defence related genes in the SL (Fig. 5, Additional file 1: Figure S7). Homologues of the *Arabidopsis* Toll-IL-1 receptor disease resistance proteins TAO1, NDR1 and RPS2 genes were up-regulated in SL. Those genes confer resistance to biotrophic bacterial pathogens in *Arabidopsis* when recognizing effectors (Coppinger *et al.*, 2004; Eitas *et al.*, 2008). In roots of beans, NDR1 also suppressed nematode parasitism by activating defence responses (McNeece *et al.*, 2017). Thus, we found indications that those proteins might also be involved in a defence response towards *P. brassicae*.

As a result of gall formation and a reduced number of fine roots, clubroot infected plants face abiotic stress like lower water and nutrient supply. The differential expression of ABA related genes (Fig. 4) are therefore likely a response to the abiotic stress in the galls. Lower water supply could also be a consequence of reduced xylem production (Malinowski *et al.*, 2012).

## Conclusions

Clubroots and symptomless roots of the same *P. brassicae* infected plants showed very different gene expression patterns. The differences in the plant hormone metabolism might be responsible for the different outcomes in gall tissue and in symptomless roots as is the increased cell wall stability in symptomless roots. These results highlight, that interpreting clubroot transcriptomes or any other data originating from whole root systems might result in a dilution of biologically relevant signatures. This clearly calls for further studies analysing intra- and inter-tissue specific pattern of clubroot infected plants. As genes involved in resistance responses to *P. brassicae* were up-regulated in symptomless roots, this might aid the identification of novel traits for resistance breeding.

## List of abbreviations

ABA: Abscisic acid
BG: Brown spindle gall
BR: Brassinosteroids
C: Control
CDS: Coding sequence
CK: Cytokinin
COG: Clusters of Orthologous Groups
DEG: Differentially expressed gene
dpi: days post inoculation
ET: Ethylene
FDR: False discovery rate
FPKM: Fragments per kilobase per million
HR: hypersensitive response
IsoPct: Isoform percentage
JA: Jasmonic acid
MeSA: Methyl-salicylate
NCBI: National Center for Biotechnology Information
ORF: Open reading frame
PbraAT: Austrian *P. brassicae* field population
PR: Pathogenesis related
RNA: Ribonucleic acid
SA: Salicylic acid
SAM: S-adenosylmethionine
SL: Symptomless root
TAIR: The Arabidopsis Information Resource
WG: White spindle gall

## Acknowledgements

We thank Srilakshmy Harikrishnan for discussion of bioinformatic analyses. M. H. Borham (Agriculture and Agri-Food Canada, 107 Science Place, Saskatoon, SK S7N0X2, Canada) provided the PT3 data. Illumina sequencing was performed at the VBCF NGS Unit (www.vbcf.ac.at).

## Availability of data and materials

The datasets generated and analysed during the current study are available in the European Nucleotide Archive (ENA; https://www.ebi.ac.uk/ena) repository under the project PRJEB26435 (Accessions ERR2567399-ERR2567408) or are available from the corresponding author on request.

## Authors’ contributions

Experimental concept and design: SN. Wet lab work: SC. Bioinformatic and statistic analysis: SC. Analysis of Results: SC, AS, SN. Manuscript writing: SC, AS, SN. Figures and tables: SC. All authors read and approved the final manuscript.

## Ethics approval and consent to participate

Not applicable.

## Consent for publication

Not applicable.

## Additional files

**Additional file 1:** Figure S1 - Figure S11

**Additional file 2:** Table S1 - Table S9

**Additional file 3:** Additional data for Figure 3

**Additional file 4:** Additional data for Figure 4

**Additional file 5:** Additional data for Figure 5

