## Supplementary Figures for "Transcriptomic response in symptomless roots of clubroot infected kohlrabi (*Brassica oleracea* var. *gongylodes*) mirrors resistant plants"

Stefan Ciaghi[1,$], Arne Schwelm[1,2,$], Sigrid Neuhauser[1,*]

*1 University of Innsbruck, Institute of Microbiology, Technikerstraße 25, 6020 Innsbruck, Austria*

*[2] Swedish University of Agricultural Sciences, Department of Plant Biology, Uppsala BioCenter, Linnean Centre for Plant Biology, P.O. Box 7080, SE-75007 Uppsala, Sweden*

[$] Contributed equally to this work

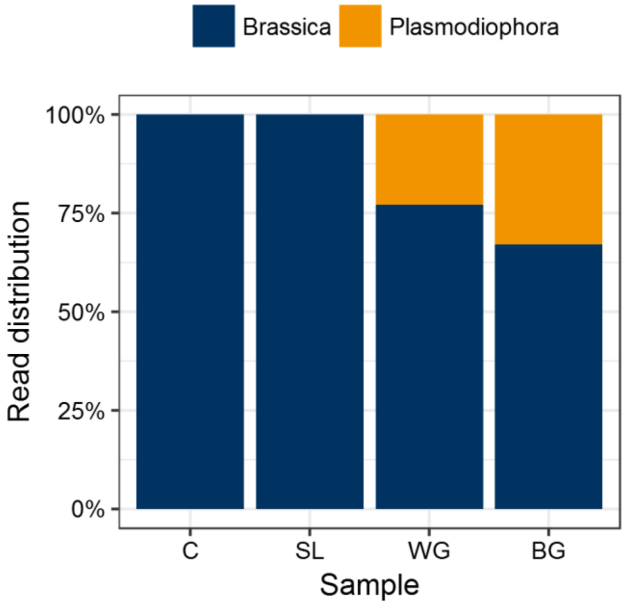


**Figure S1: Kohlrabi (*Brassica oleracea*) and *Plasmodiophora brassicae* reads.** Samples from the control plant (C) and symptomless roots (SL) of infected plants contained less than 0.0005% *P. brassicae* reads. *P. brassicae* read partition increased from 23% in WG to 33% in BG.

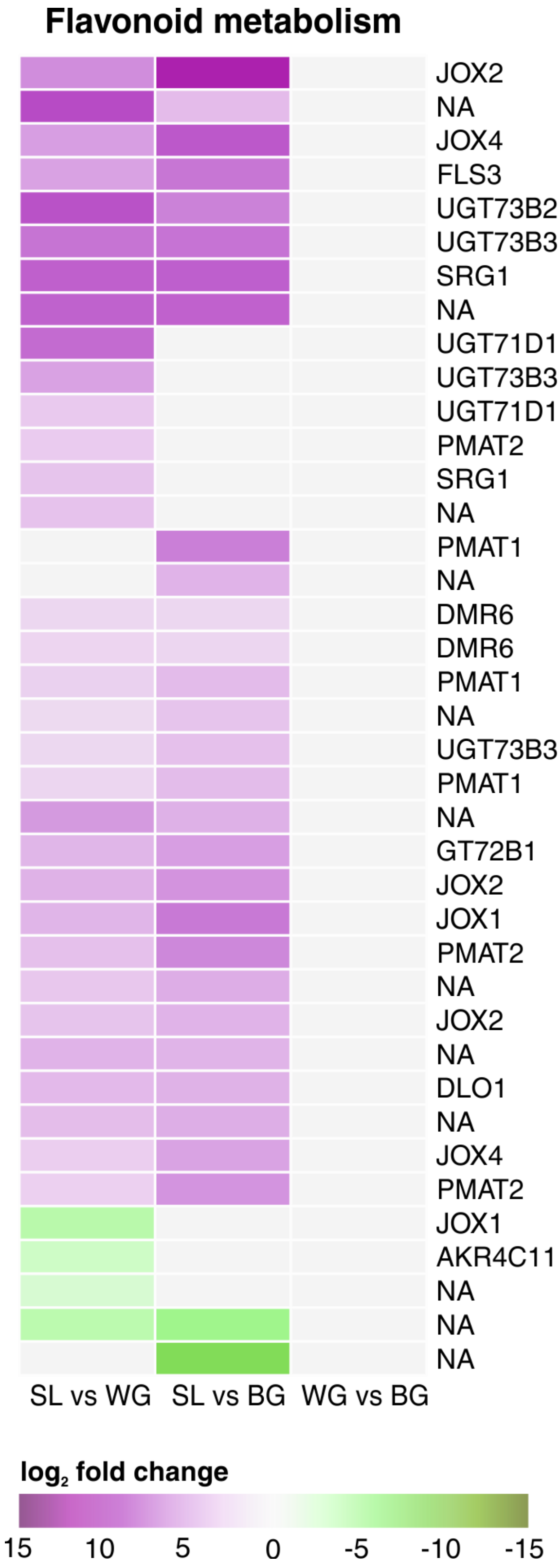


**Figure S2: Kohlrabi flavonoid metabolism.** Clustered heatmaps of $\log_2$ fold change values of DEGs. No DEGs were present comparing WG with BG. Up-regulated genes are shaded in purple and down-regulated genes in green. *Arabidopsis* homologs are given. NA: not assigned.

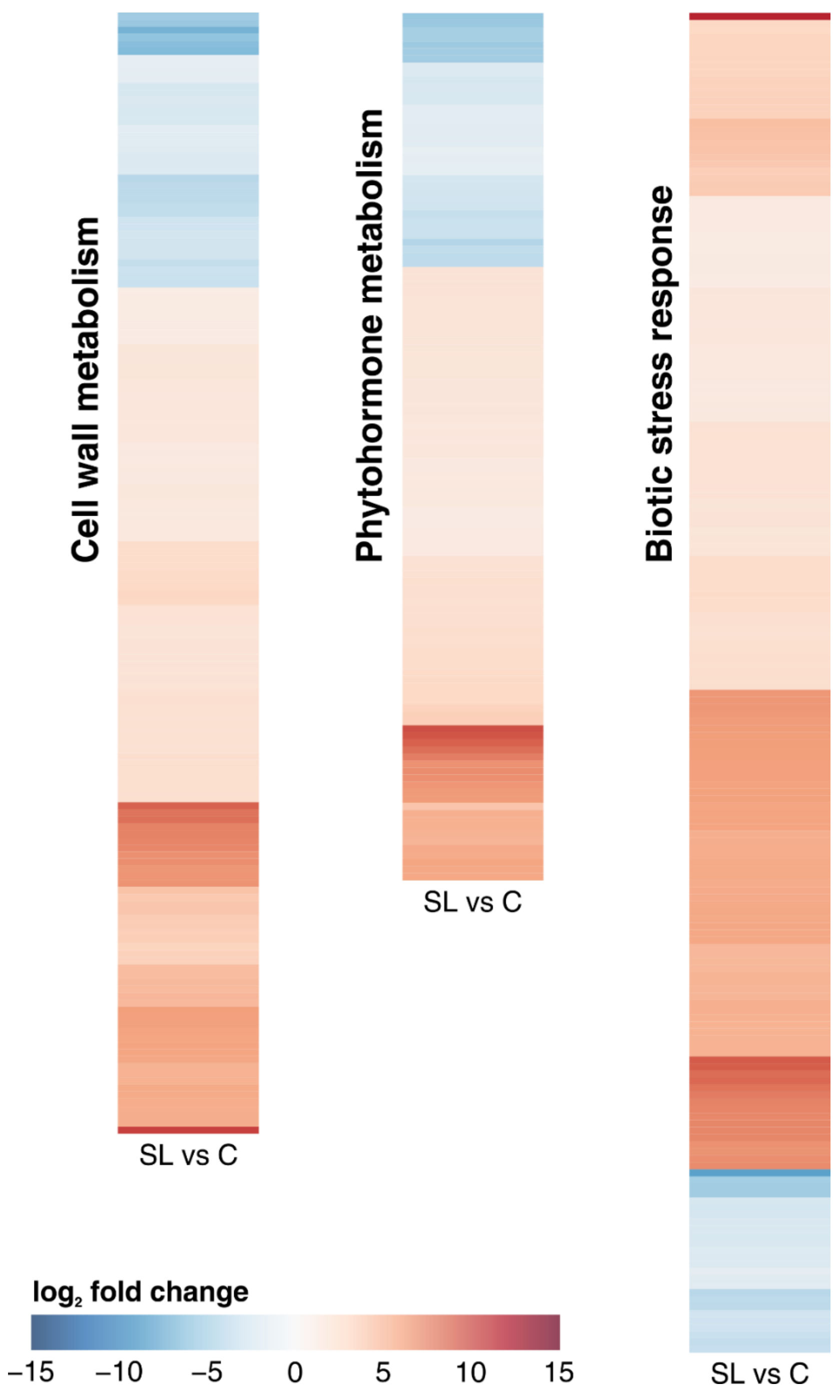


**Figure S3: Kohlrabi DEGs comparing symptomless roots (SL) of infected plants with the uninfected control plant (C).** Heatmaps are given for cell wall metabolism, phytohormone metabolism, and biotic stress response. Up-regulated genes are shaded in red and down-regulated genes in blue. Gene identifiers are not illustrated due to readability.

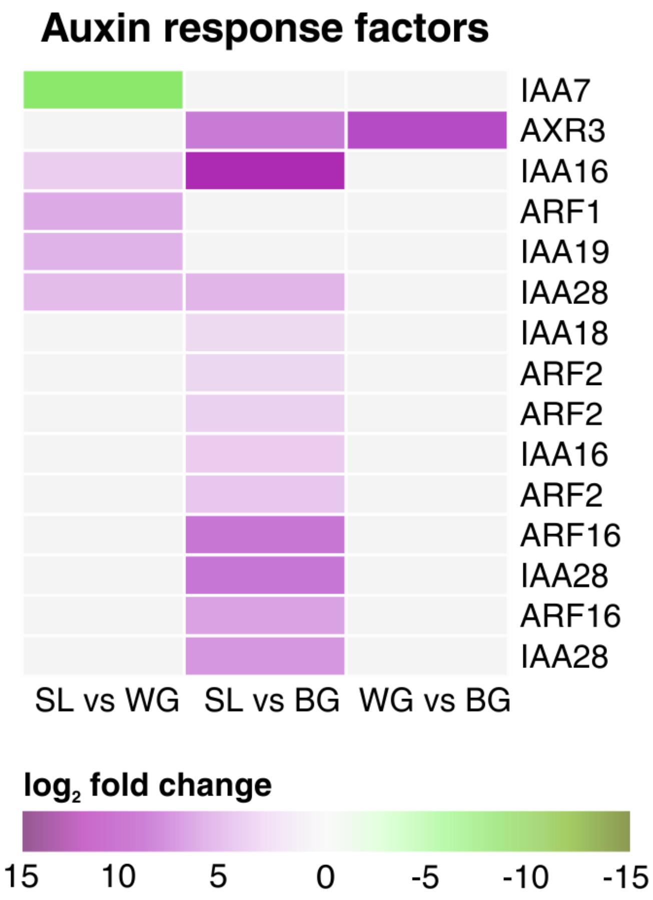


**Figure S4: Kohlrabi auxin response factors.** Clustered heatmaps of $log_2$ fold change values of DEGs. One DEG (AXR3) was present comparing WG with BG. Up-regulated genes are shaded in purple and down-regulated genes in green. *Arabidopsis* homologs are given.

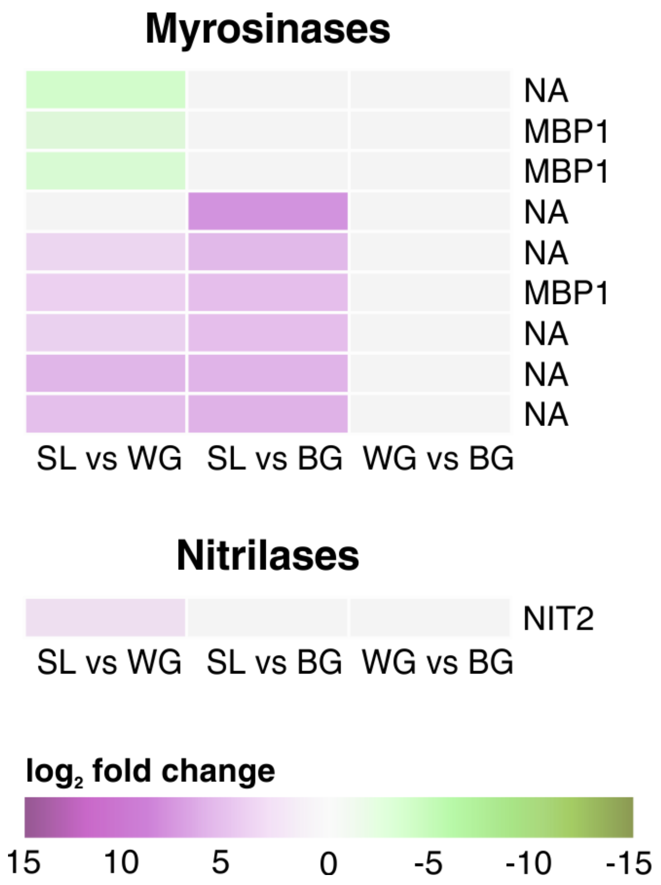


**Figure S6: Kohlrabi myrosinases and nitrilases.** Clustered heatmaps of $log_2$ fold change values of DEGs. No DEGs were present comparing WG with BG. Up-regulated genes are shaded in purple and down-regulated genes in green. *Arabidopsis* homologs are given. NA: not assigned.

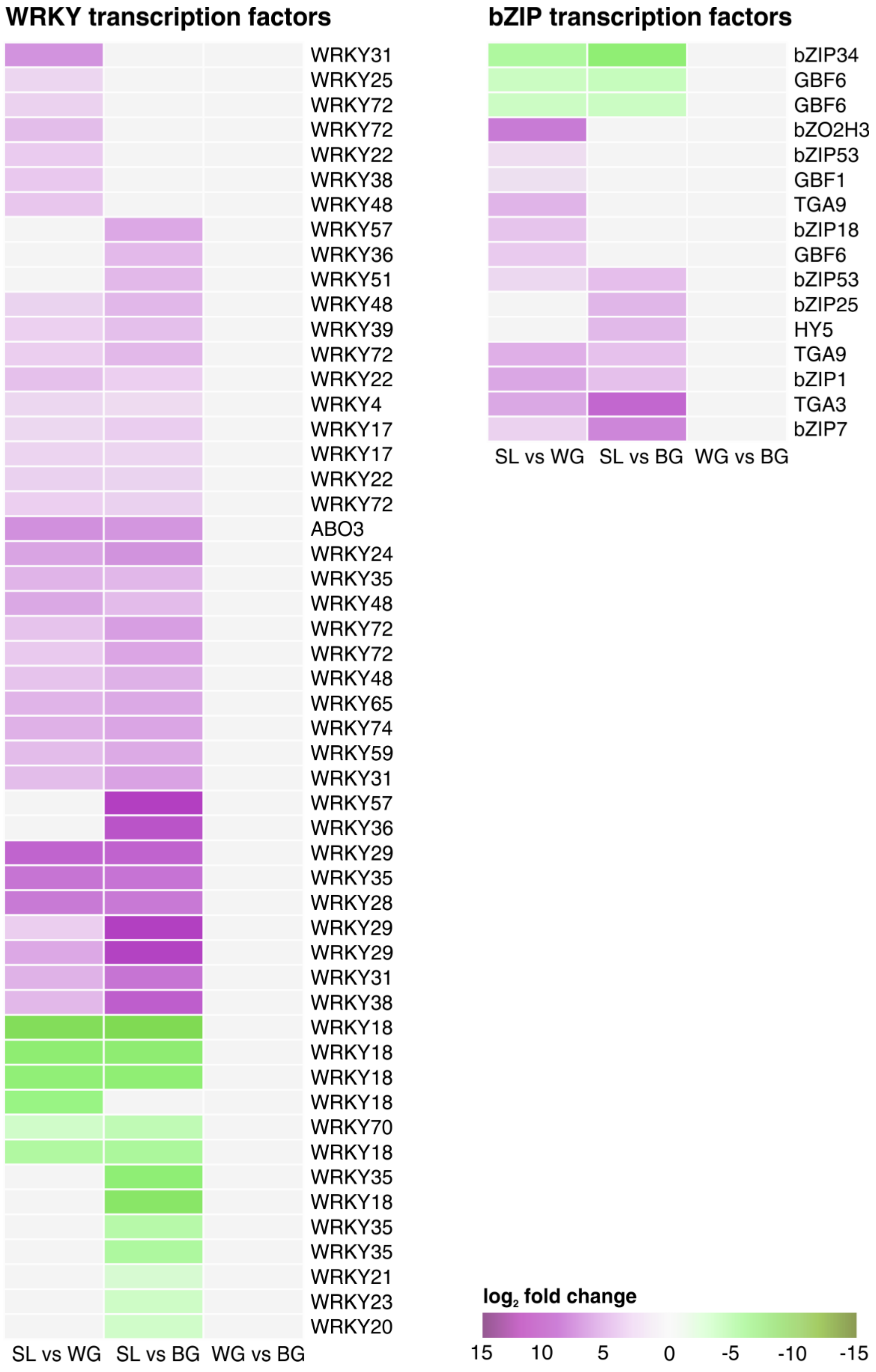


**Figure S6: Kohlrabi WRKY and bZIP transcription factors.** Clustered heatmaps of $\log_2$ fold change values of DEGs. No DEGs were present comparing WG with BG. Up-regulated genes are shaded in purple and down-regulated genes in green. *Arabidopsis* homologs are given. NA: not assigned.

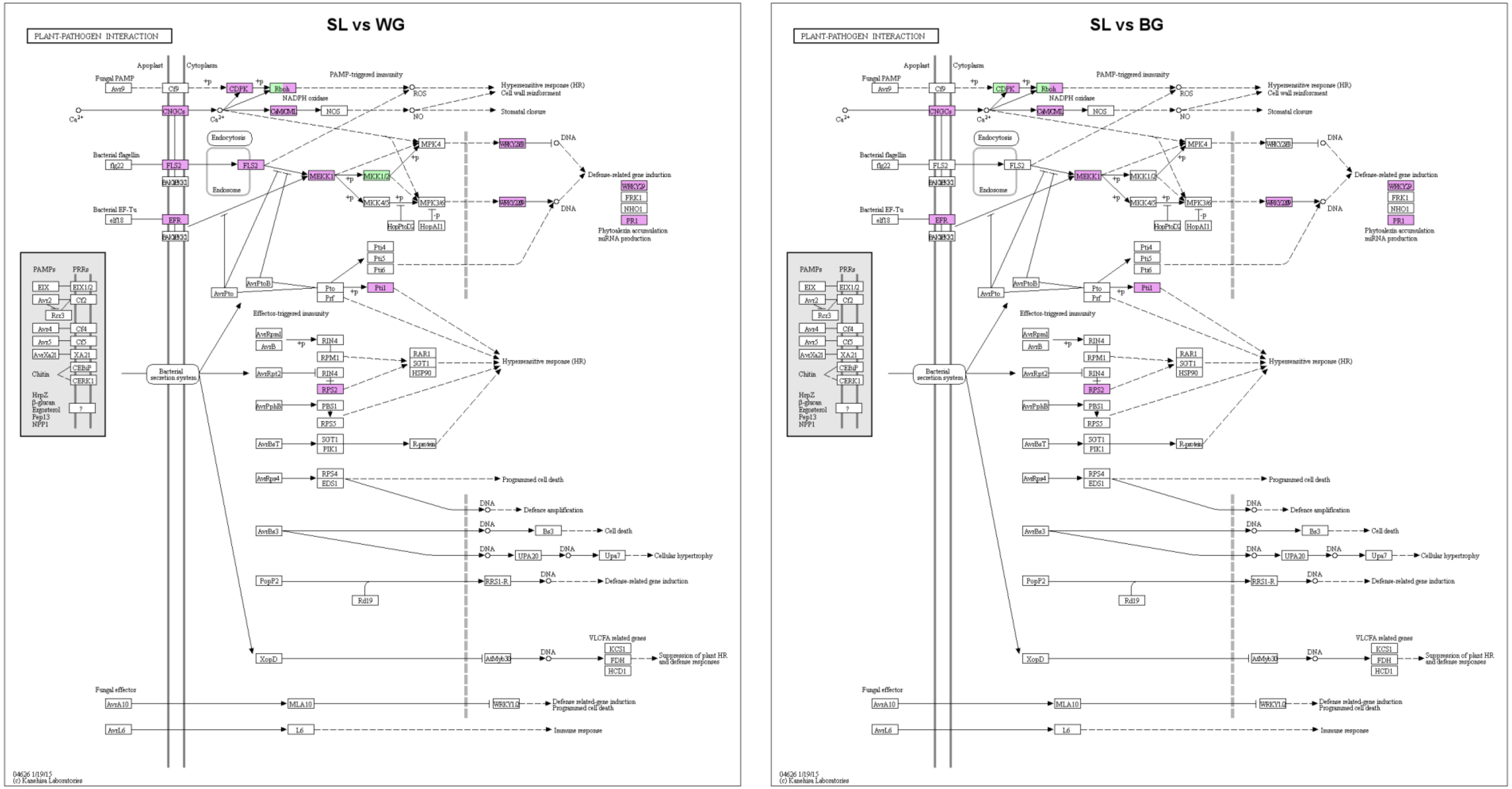


**Figure S7: KEGG map for plant-pathogen interaction.** Up-regulated genes in SL compared with galls are shaded in purple and down-regulated genes in green. Genes shaded in purple and green were up- and down-regulated in different homologs or isoforms.

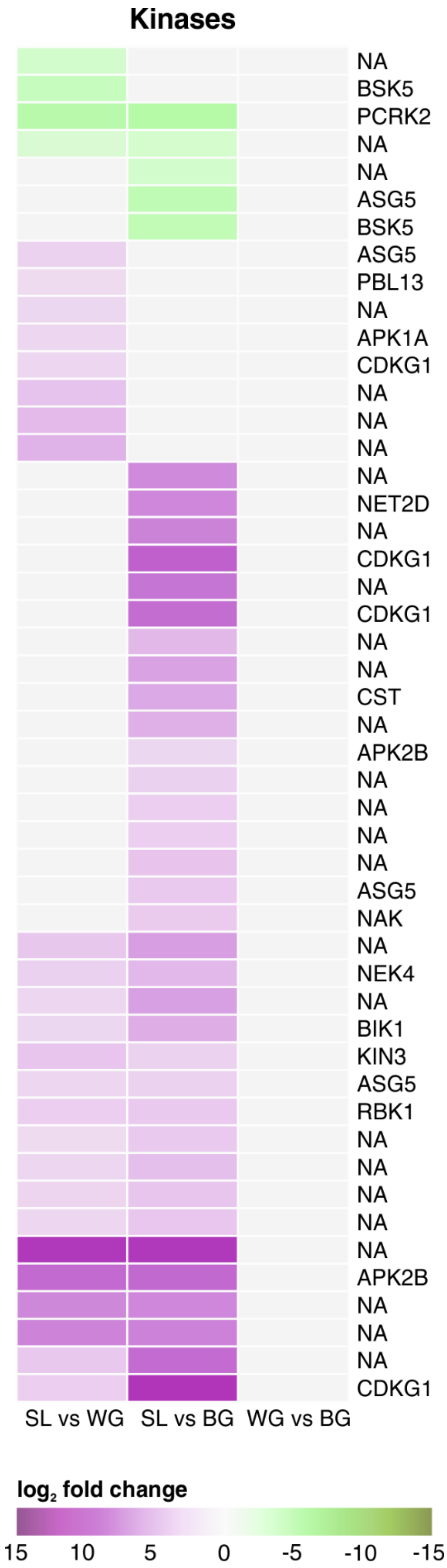


**Figure S8: Kohlrabi kinases.** Clustered heatmaps of $\log_2$ fold change values of DEGs. No DEGs were present comparing WG with BG. Up-regulated genes are shaded in purple and down-regulated genes in green. *Arabidopsis* homologs are given. NA: not assigned.

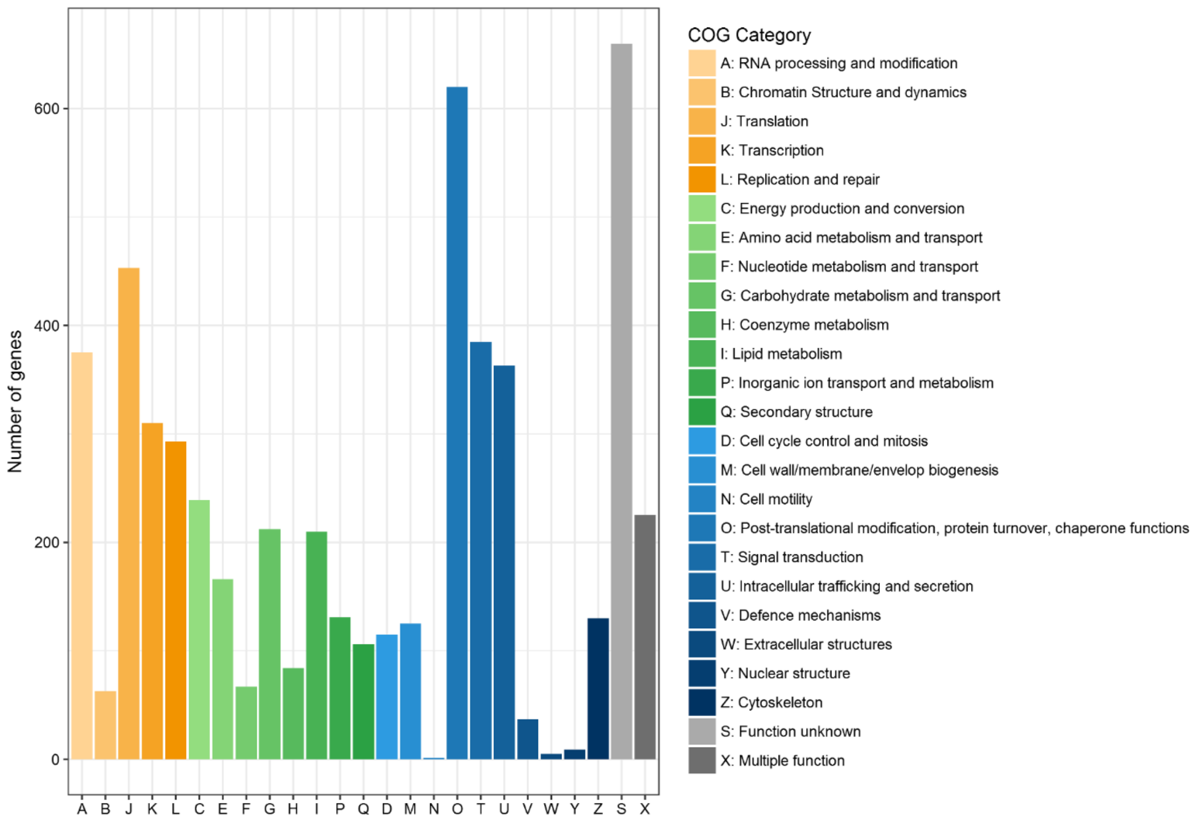


**Figure S9: Numbers of *P. brassicae* genes in clubroot infected kohlrabi roots per COG category.** Bars indicate total genes found across all libraries. Unassigned genes (n = 5482) are not illustrated. Orange: Information and Storage Processing; Green: Metabolism; Blue: Cellular Process and Signalling; Grey: Poorly Characterized.

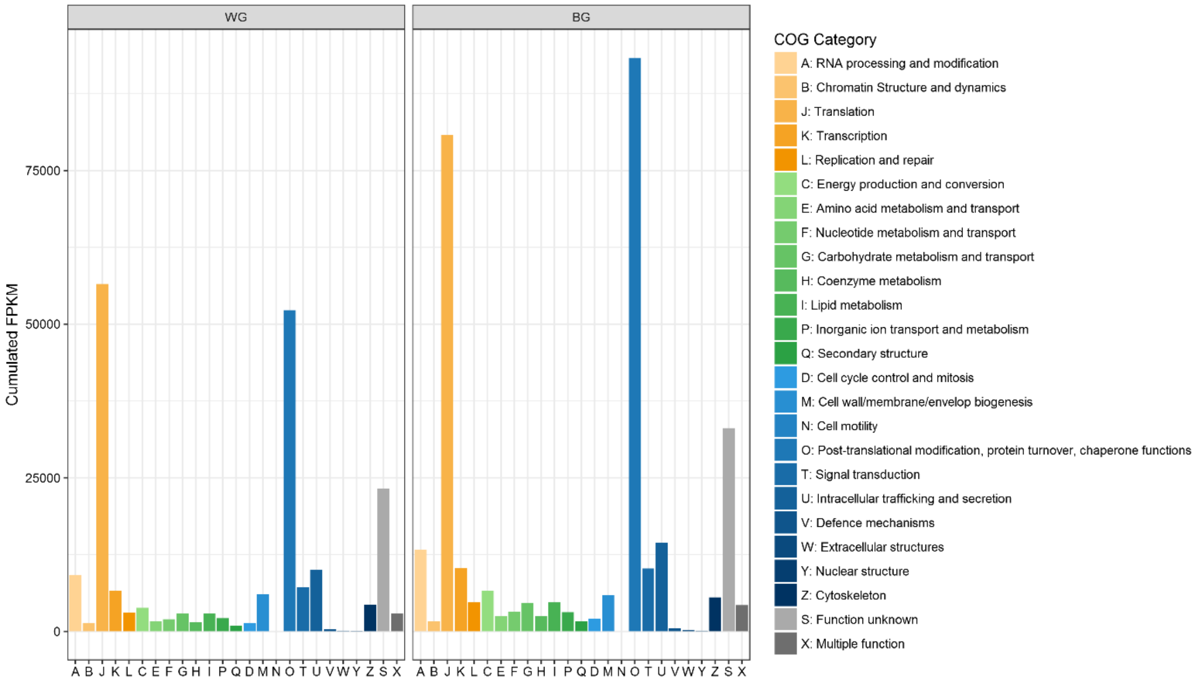


**Figure S10: Cumulated FPKM values of *P. brassicae* reads obtained from WG and BG samples.** Unassigned genes are not illustrated. Orange: Information and Storage Processing; Green: Metabolism; Blue: Cellular Process and Signalling; Grey: Poorly Characterized.

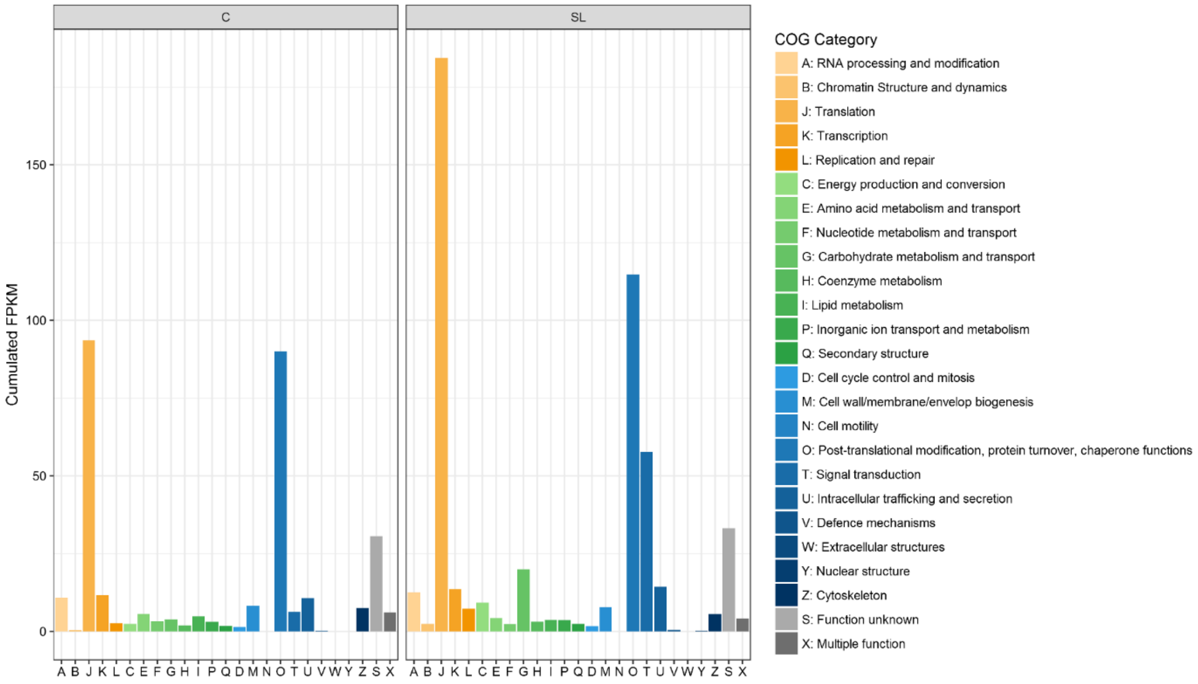


**Figure S11: Cumulated FPKM values of *P. brassicae* reads obtained from C and SL samples.** Unassigned genes are not illustrated. Orange: Information and Storage Processing; Green: Metabolism; Blue: Cellular Process and Signalling; Grey: Poorly Characterized.
