## Supplementary Tables for "Transcriptomic response in symptomless roots of clubroot infected kohlrabi (*Brassica oleracea* var. *gongylodes*) mirrors resistant plants"

Stefan Ciaghi[1,$], Arne Schwelm[1,2,$], Sigrid Neuhauser[1,*]

[1] *University of Innsbruck, Institute of Microbiology, Technikerstraße 25, 6020 Innsbruck, Austria*

[2] *Swedish University of Agricultural Sciences, Department of Plant Biology, Uppsala BioCenter, Linnean Centre for Plant Biology, P.O. Box 7080, SE-75007 Uppsala, Sweden*

[$] Contributed equally to this work

**Table S1: Total (raw) and good quality reads of the sequenced libraries.** Only reads with a minimum length of 75 bp and a sliding window of 5 bp and an average quality score > 20 were kept. Illumina adapters were removed.

| Library | Total reads | Good quality reads |
|---|---:|---:|
| C | 35470516 | 18724257 |
| SL-1 | 25749647 | 17112091 |
| SL-2 | 26086772 | 15292763 |
| SL-3 | 22998692 | 13487038 |
| WG-1 | 23140251 | 15398974 |
| WG-2 | 21623635 | 13055601 |
| WG-3 | 30159869 | 16748310 |
| BG-1 | 27662722 | 15482391 |
| BG-2 | 33728086 | 17655530 |
| BG-3 | 31782654 | 19236687 |
| **Total** | **278402844** | **162193642** |

**Table S2: Kohlrabi DEGs compared across the three root tissue types.**

| Type of DEGs | SL vs WG | SL vs BG | WG vs BG |
|---|---|---|---|
| Up-regulated DEGs | 1619 | 942 | 6 |
| Down-regulated DEGs | 2280 | 2571 | 13 |
| Total DEGs | 3899 | 3513 | 19 |

**Table S3: Kohlrabi DEGs per COG category in infected plants.** Total DEGs of SL vs WG, SL vs BG, and WG vs BG are given. Individual DEGs can occur in more than one comparison but are counted only once. The five categories containing most DEGs are highlighted in bold. Orange: Information and Storage Processing; Green: Metabolism; Blue: Cellular Process and Signalling; Grey: Poorly Characterized.

| COG category | # of DEGs |
|---|---|
| A: RNA processing and modification | 74 |
| B: Chromatin structure and dynamics | 30 |
| C: Energy production and conservation | 130 |
| D: Cell cycle control and mitosis | 87 |
| E: Amino acid metabolism and transport | 179 |
| F: Nucleotide metabolism and transport | 42 |
| **G: Carbohydrate metabolism and transport** | **388** |
| H: Coenzyme metabolism | 46 |
| I: Lipid metabolism | 170 |
| J: Translation | 136 |
| **K: Transcription** | **479** |
| L: Replication and repair | 71 |
| M: Cell wall/membrane/envelop biogenesis | 52 |
| N: Cell motility | 0 |
| **O: Post-translational modification, protein turnover, chaperon functions** | **349** |
| P: Inorganic ion transport and metabolism | 195 |
| Q: Secondary Structure | 251 |
| R: General function prediction only | 0 |
| **S: Function unknown** | **1435** |
| **T: Signal transduction** | **538** |
| U: Intracellular trafficking and secretion | 136 |
| V: Defence mechanisms | 30 |
| W: Extracellular structures | 3 |
| X: Multiple function | 127 |
| Y: Nuclear structure | 1 |
| Z: Cytoskeleton | 49 |
| N/A: Unassigned | 206 |

**Table S4: Complete list of kohlrabi DEGs between BG and WG.** The Functional annotations are given as returned by eggNOG mapper. Up-regulated genes are highlighted in red, down-regulated genes in blue.

| Transcript ID | Functional annotation | $log_2$ fold change |
|---|---|---|
| Bol 94671.3.1.4 | F-box kelch-repeat protein | 10.18 |
| Bol 99971.0.1.7 | Acyl-CoA synthetase long-chain family | 8.88 |
| Bol 76700.1.2.2 | chitinase | 8.72 |
| Bol 97623.1.1.1 | Aspartyl protease family protein | 8.19 |
| Bol 94026.0.2.1 | GDE1 homolog | 8.13 |
| Bol 159044.0.1.1 | inhibitor | 5.20 |
| Bol 99469.1.6.8 | Glutamate decarboxylase | -3.77 |
| Bol 93718.3.1.2 | Indole-3-acetic acid-amido synthetase | -6.15 |
| Bol 134791.0.1.1 | expressed protein | -7.21 |
| Bol 100433.0.1.1 | Armadillo/beta-catenin-like repeat | -8.09 |
| Bol 85918.1.1.4 | May negatively regulate the SNF1 kinase | -8.15 |
| Bol 96726.0.1.1 | Farnesyl-diphosphate farnesyltransferase | -8.24 |
| Bol 99205.7.2.8 | NA | -8.36 |
| Bol 96830.0.1.6 | phosphatase 2C | -8.38 |
| Bol 96394.0.1.1 | glycosyltransferase family 14 protein | -8.40 |
| Bol 98490.1.1.3 | RNA recognition motif containing protein | -8.53 |
| Bol 96780.1.1.3 | AUX/IAA family | -8.57 |
| Bol 99251.0.2.4 | Cytochrome P450 | -8.62 |
| Bol 64904.0.1.1 | Plastocyanin-like domain | -10.61 |

**Table S5: DEGs between the uninfected control plant and the three tissue types of infected kohlrabi plants (SL, WG, BG).**

| Type of DEGs | SL vs C | WG vs C | BG vs C |
|---|---|---|---|
| Up-regulated DEGs | 4606 | 8347 | 6031 |
| Down-regulated DEGs | 1721 | 5075 | 5390 |
| Total DEGs | 6327 | 13422 | 11421 |

**Table S6: Kohlrabi DEGs per COG category between infected and control plants.** Total DEGs of SL vs C, WG vs C, and BG vs C are given. Individual DEGs can occur in more than one comparison but are counted only once. The five categories containing most DEGs are highlighted in bold. Orange: Information and Storage Processing; Green: Metabolism; Blue: Cellular Process and Signalling; Grey: Poorly Characterized.

| COG category | # of DEGs |
|---|---|
| A: RNA processing and modification | 504 |
| B: Chromatin structure and dynamics | 141 |
| C: Energy production and conservation | 469 |
| D: Cell cycle control and mitosis | 289 |
| E: Amino acid metabolism and transport | 528 |
| F: Nucleotide metabolism and transport | 211 |
| G: Carbohydrate metabolism and transport | 1149 |
| H: Coenzyme metabolism | 203 |
| I: Lipid metabolism | 496 |
| J: Translation | 681 |
| **K: Transcription** | **1636** |
| L: Replication and repair | 403 |
| M: Cell wall/membrane/envelop biogenesis | 182 |
| N: Cell motility | 0 |
| **O: Post-translational modification, protein turnover, chaperon functions** | **1566** |
| P: Inorganic ion transport and metabolism | 605 |
| Q: Secondary Structure | 563 |
| R: General function prediction only | 0 |
| **S: Function unknown** | **5081** |
| **T: Signal transduction** | **1830** |
| U: Intracellular trafficking and secretion | 672 |
| V: Defence mechanisms | 102 |
| W: Extracellular structures | 11 |
| X: Multiple function | 528 |
| Y: Nuclear structure | 2 |
| Z: Cytoskeleton | 223 |
| **N/A: Unassigned** | **1155** |

**Table S7: Twenty highest expressed genes of *Plasmodiophora brassicae* in WG.** Annotation given as predicted with eggNOG mapper. Putative secreted proteins are tagged (●). FPKM: fragments per kilobase (of exons) per million reads. NA: not assigned.

| Transcript ID | Functional annotation | FPKM | Secreted |
|---|---|---|---|
| PbraAT 83084.17.1.2 | NA | 14838 | |
| PbraAT 133546.1.1.1 | glutathione-S-transferase | 12890 | |
| PbraAT 108093.0.1.1 | NA | 10438 | |
| PbraAT 146283.0.1.1 | heat shock protein | 9304 | |
| PbraAT 83084.17.1.1 | NA | 7262 | |
| PbraAT 83084.18.1.1 | NA | 7031 | ● |
| PbraAT 158899.8.1.1 | NA | 6733 | |
| PbraAT 100779.1.1.1 | PPIases accelerate the folding of proteins | 5484 | |
| PbraAT 133446.2.1.1 | carboxyl methyltransferase **(PbBSMT)** | 4983 | ● |
| PbraAT 120659.1.1.1 | ribosomal protein | 3330 | |
| PbraAT 108093.5.1.1 | ankyrin repeat domain-containing protein | 3168 | |
| PbraAT 108221.0.1.1 | 60s ribosomal protein | 3167 | |
| PbraAT 108279.2.1.1 | ribosomal protein | 3084 | |
| PbraAT 108037.1.1.1 | finger protein | 3048 | |
| PbraAT 100599.0.1.1 | ribosomal protein | 2332 | |
| PbraAT 158960.0.1.1 | 60s ribosomal protein | 2271 | |
| PbraAT 158883.0.1.1 | NA | 2113 | |
| PbraAT 158922.0.1.1 | NA | 1789 | |
| PbraAT 121753.0.1.1 | ribosomal protein L12 | 1620 | |
| PbraAT 146514.1.1.1 | NA | 1561 | |

**Table S8: Twenty highest expressed genes of *Plasmodiophora brassicae* in BG.** Annotation given as predicted with eggNOG mapper. Putative secreted proteins are tagged (●). FPKM: fragments per kilobase (of exons) per million reads. NA: not assigned.

| Transcript ID | Functional annotation | FPKM | Secreted |
|---|---|---|---|
| PbraAT 133546.1.1.1 | glutathione-S-transferase | 25036 | |
| PbraAT 146283.0.1.1 | heat shock protein | 18301 | |
| PbraAT 83084.17.1.2 | NA | 12855 | |
| PbraAT 100779.1.1.1 | PPIases accelerate the folding of proteins | 8838 | |
| PbraAT 158899.8.1.1 | NA | 8751 | |
| PbraAT 108093.0.1.1 | NA | 8468 | |
| PbraAT 133446.2.1.1 | carboxyl methyltransferase **(PbBSMT)** | 7182 | ● |
| PbraAT 83084.17.1.1 | NA | 6570 | |
| PbraAT 83084.18.1.1 | NA | 6020 | ● |
| PbraAT 146514.1.1.1 | NA | 5713 | |
| PbraAT 120659.1.1.1 | ribosomal protein | 4744 | |
| PbraAT 108279.2.1.1 | ribosomal protein | 3716 | |
| PbraAT 133396.0.1.1 | heat shock protein | 3528 | |
| PbraAT 100599.0.1.1 | ribosomal protein | 3503 | |
| PbraAT 108221.0.1.1 | 60s ribosomal protein | 3366 | |
| PbraAT 158960.0.1.1 | 60s ribosomal protein | 3170 | |
| PbraAT 146422.1.1.1 | peroxiredoxin | 2950 | |
| PbraAT 158936.0.1.1 | 10 kda heat shock protein | 2929 | |
| PbraAT 158883.0.1.1 | NA | 2630 | |
| PbraAT 161208.0.1.1 | ribosomal protein | 2604 | |

**Table S9: *Plasmodiophora brassicae* DEGs between BG and WG.** Annotations are given as predicted with eggNOG mapper. Up-regulated genes in BG are highlighted in red, down-regulated genes in BG in blue.

| Transcript ID | Functional annotation | $\log_2$ fold change |
|---|---|---|
| PbraAT 71424.0.1.5 | heat shock protein | 11.45 |
| PbraAT 52700.0.2.1 | DNA-directed RNA polymerase | 9.56 |
| PbraAT 26420.0.2.1 | structural maintenance of chromosomes | 9.40 |
| PbraAT 69953.0.1.2 | Scl Tal1 interrupting locus | 8.82 |
| PbraAT 93256.0.5.1 | retrotransposon protein | -8.36 |
